# Epigenetic Interaction between UTX and DNMT1 Regulates Diet-Induced Myogenic Remodeling in Brown Fat

**DOI:** 10.1101/2020.08.05.238923

**Authors:** Fenfen Li, Jia Jing, Miranda Movahed, Xin Cui, Qiang Cao, Rui Wu, Ziyue Chen, Liqing Yu, Yi Pan, Huidong Shi, Bingzhong Xue, Hang Shi

**Author notes:** Correspondence should be addressed to: Hang Shi, Department of Biology, Georgia State University, Atlanta, GA 30303, USA. Contact. Bingzhong Xue, Department of Biology, Georgia State University, Atlanta, GA 30303, USA. Contact.

## Abstract

Brown adipocytes share the same developmental origin with skeletal muscle. Here we find that a brown adipocyte-to-myocyte remodeling also exists in mature brown adipocytes, and is induced by prolonged high fat diet (HFD) feeding, leading to brown fat dysfunction. This process is regulated by the interaction of epigenetic pathways involving histone and DNA methylation. In mature brown adipocytes, the histone demethylase UTX maintains persistent demethylation of the repressive mark H3K27me3 at *Prdm16* promoter, leading to high *Prdm16* expression. PRDM16 then recruits DNA methyltransferase DNMT1 to *Myod1* promoter, causing *Myod1* promoter hypermethylation and suppressing its expression. The interaction between PRDM16 and DNMT1 coordinately serves to maintain brown adipocyte identity while repressing myogenic remodeling in mature brown adipocytes, thus promoting their active brown adipocyte thermogenic function. Suppressing this interaction by HFD feeding induces brown adipocyte-to-myocyte remodeling, which limits brown adipocyte thermogenic capacity and compromises diet-induced thermogenesis, leading to the development of obesity.

## Introduction

Obesity is now considered as an epidemic disorder that poses as an independent risk factor for the development of various metabolic disorders such as insulin resistance/type 2 diabetes, hypertension, dyslipidemia, and cardiovascular diseases (Hill et al., 2012). Chronic energy excess due to energy intake over energy expenditure results in obesity (Hill et al., 2012). Thus, understanding the mechanism underlying the regulation of energy homeostasis may help identify the therapeutic targets for the prevention and treatment of obesity.

Total energy expenditure consists of three aspects: energy required for basal metabolic rate, energy expended to perform physical activity and energy used to generate heat. The last one is termed as adaptive thermogenesis, which mainly takes place in brown adipose tissue (BAT) due to the unique presence of uncoupling protein 1 (UCP1) in the inner membrane of mitochondria (Donahoo et al., 2004; Hill et al., 2012). UCP1 acts to uncouple oxidative phosphorylation from ATP synthesis, thereby dissipating energy as heat and profoundly increasing overall energy expenditure (Cannon and Nedergaard, 1985; Nicholls and Locke, 1984). In addition, recent studies have reported the existence of UCP1-independent thermogenesis that is generated by sarco/endoplasmic reticulum Ca^2+^-ATPase 2b/ATPase, Ca^2+^ transporting, cardiac muscle, slow twitch 2 (SERCA2b/ATP2a2)-mediated calcium cycling or creatine-driven substrate cycling (Ikeda et al., 2017; Kazak et al., 2015). Given the presence of human brown fat that profoundly increases energy expenditure (Cypess et al., 2009; van Marken Lichtenbelt et al., 2009; Virtanen et al., 2009), better understanding the mechanism underlying BAT thermogenesis has a translational implication for the treatment of obesity.

Most of the current studies investigating the mechanisms in the regulation of brown fat thermogenesis focus on cellular signaling pathways; much less is known about the epigenetic mechanisms in this process. We have recently discovered several epigenetic pathways involved in adipocyte development and brown adipocyte thermogenesis (Chen et al., 2016; Li et al., 2016; Yang et al., 2016; Zha et al., 2015). For instance, we have reported that ubiquitously transcribed tetratricopeptide/ lysine (K)-specific demethylase 6A (*Utx/Kdm6a*), a histone demethylase that preferentially catalyzes the demethylation of tri-methylated histone H3 lysine 27 (H3K27me3) and therefore relieves its ability of gene silencing (Ge, 2012), plays a key role in regulating brown adipocyte thermogenic program via a coordinated regulation of H3K27 demethylation and acetylation (Zha et al., 2015). Specifically, *Utx*, whose expression in white adipose tissue (WAT) or BAT is induced by cold exposure, acts as a positive regulator of BAT thermogenic gene expression (Zha et al., 2015). However, the physiological significance of *Utx* in the regulation of energy homeostasis remains unknown. In this study, we have generated mice with brown adipocyte-specific *Utx* knockout (UTXKO) and characterized metabolic phenotypes of these mice. We have further interrogated potential epigenetic mechanisms underlying *Utx*’s regulation of brown fat function during diet-induced obesity (DIO), and found that this process involves an interaction between histone and DNA methylation in the promoters of key molecules regulating brown or myogenic lineage determination, leading to a myogenic remodeling and thermogenic dysfunction in BAT of UTXKO mice during the development of DIO. We have also identified that BAT-to-myocyte remodeling in BAT represents a common feature in DIO, which eventually leads to BAT dysfunction and contribute to the development of DIO.

## Results

### Mice with UTX deficiency in brown fat exhibit impaired BAT thermogenesis and are susceptible to DIO

We previously reported that *Utx* knockdown reduces mRNA levels of brown-specific genes, whereas overexpression of *Utx* does the opposite in cultured brown adipocyte cell lines (Zha et al., 2015). However, it remains unknown whether *Utx* regulates BAT thermogenic function and whole-body energy homeostasis *in vivo*. In the current study, we first measured *Utx* expression pattern in brown and white adipose tissues. *Utx* mRNA level was higher in interscapular brown adipose tissue (iBAT) than in inguinal white adipose tissue (iWAT) and epididymal WAT (eWAT), and was induced in both iBAT and iWAT by a 7-day 5°C cold challenge (**Suppl. Fig 1A** and **1B**).

To determine the role of *Utx* in the regulation of BAT thermogenic function and energy homeostasis *in vivo*, we generated mice (UTXKO) with specific deletion of *Utx* in brown fat by crossing *Utx*-floxed mice (Welstead et al., 2012) with *Ucp1*-cre mice (Kong et al., 2014). *Utx* is located on the X chromosome but escapes X chromosome inactivation in females (Greenfield et al., 1998). Thus, female UTXKO mice were defined as homozygous *Utx*^fl/fl^ with *Ucp1-Cre* (*Ucp1-Cre::Utx^fl/fl^*), with *Utx^fl/fl^* littermates as control; whereas male UTXKO mice were defined as hemizygous *Utx^fl/Y^* with *Ucp1-Cre* (*Ucp1-Cre::Utx^fl/Y^*), with *Utx^fl/Y^* littermates as control. Male and female UTXKO and their fl/Y or fl/fl littermate control mice were viable and were born with expected Mendelian frequency, and as expected, *Utx* deletion in brown fat resulted in an 80% reduction of *Utx* mRNA expression in male UTXKO mice compared to their fl/Y littermates (**Suppl. Fig 2A**).

Although there was no difference in body weight of male UTXKO mice compared to that of fl/Y mice fed a regular chow diet (**Suppl. Fig 2B**), UTXKO mice exhibited a significant increase in fat mass and a significant decrease in lean mass (**Suppl. Fig 2C**) measured by a Minispec NMR body composition analyzer. In consistence, we also found significantly increased individual fat pad mass including iBAT and iWAT, and a tendency of increased eWAT and retroperitoneal WAT (rWAT) mass in male UTXKO mice (**Supp. Fig 2D**). This was associated with larger adipocytes in iBAT and iWAT (**Suppl. Fig 2E**) and less UCP1 immunohistochemistry (IHC) staining in iBAT (**Suppl. Fig 2F**), suggesting a potential decrease in brown fat thermogenesis in brown fat of UTXKO mice compared to their fl/Y littermates. With increased adiposity, male UTXKO mice exhibited an increase in fed and fast glucose levels and fed insulin levels (**Suppl. Fig 2G**), suggesting that these UTXKO mice had impaired insulin sensitivity compared to their fl/Y littermate controls. Indeed, this was further confirmed by impaired glucose tolerance test (GTT) (**Suppl. Fig 2H**) with increased circulating insulin levels at 15 minutes during GTT in male UTXKO mice compared to that of fl/Y mice (**Suppl. Fig 2I**).

To determine the role of brown fat UTX in the regulation of diet-induced obesity, we conducted metabolic characterization of body weight, energy metabolism and insulin sensitivity in UTXKO mice fed a high fat diet (HFD). As shown in **Fig 1A**, male UTXKO mice gained significantly more weight compared to their fl/Y littermates on the HFD. Using a Minispec NMR body composition analyzer, we found that UTXKO mice exhibited a significant increase in fat mass and a significant decrease in lean mass (**Fig 1B**). In consistence, iWAT and iBAT mass was also significantly increased in UTXKO mice compared to their fl/Y littermates (**Fig 1C**) with larger adipocytes in iBAT, iWAT and eWAT (**Fig 1D**). Using a PhenoMaster metabolic cage system, we found that UTXKO mice displayed lower energy expenditure and oxygen consumption (**Fig 1E** and **Suppl. Fig 3A**), whereas there was no difference in locomotor activity, respiratory exchange rate (RER) and food intake between UTXKO and fl/Y mice (**Suppl. Fig 3B-D**). These data indicate that reduced energy expenditure may primarily account for the obese phenotype in UTXKO mice fed the HFD. Quantitative RT-PCR analysis revealed significantly reduced thermogenic gene expression, including *Ucp1*, *Prdm16*, *Pgc1α*, *Pgc1β*, *Ebf2*, *Ebf3*, *Cpt1b*, *Cox1*, *Cidea*, *Pparγ*. *Dio2* and *Otop1* (**Fig 1F**) in iBAT of UTXKO mice, which was associated with decreased UCP1 protein levels as measured by both immunoblotting and UCP1 IHC staining (**Fig 1G** and **1H**). UTXKO mice also displayed glucose intolerance and insulin resistance as assessed by glucose and insulin tolerance tests (GTT and ITT, respectively) (**Fig 1I** and **1J**). These data indicate that mice with *Utx* deficiency in brown fat have increased adiposity with impaired insulin sensitivity when fed a regular chow diet, and are susceptible to diet-induced obesity with exacerbated insulin resistance when fed HFD.

**Figure 1.**
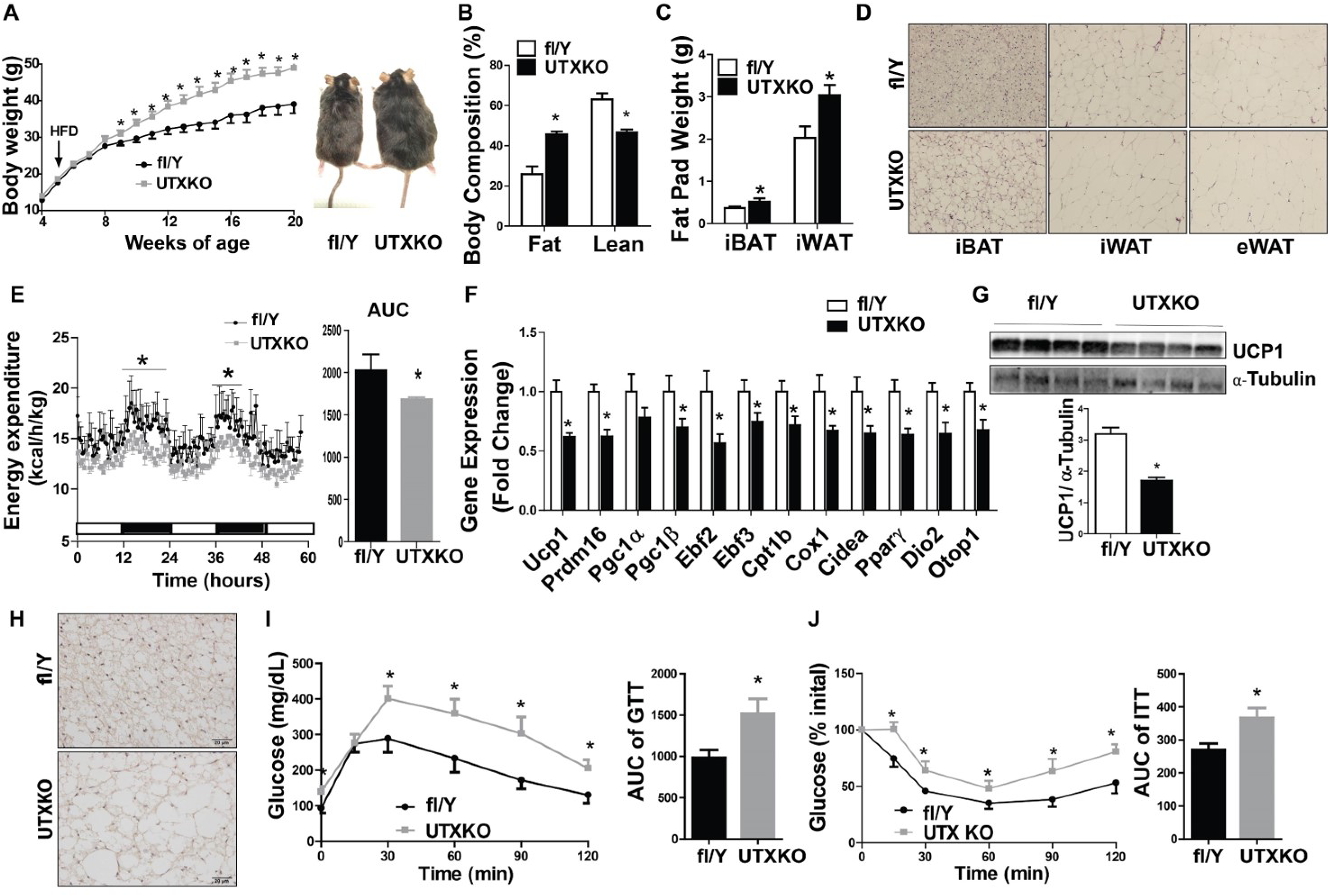
UTX deficiency in brown fat promotes high fat diet (HFD)-induced obesity. Male UTXKO and their littermate control fl/Y mice were put on HFD when they were 5 weeks of age. (A)-(C) Body weight growth curve (A), Body composition (B), and Fat pad weight (interscapular brown adipose tissue (iBAT), inguinal white adipose tissue (iWAT) and epididymal WAT (eWAT)) (C) in male UTXKO and fl/Y mice fed HFD (n=6/group). (D) H&E staining of iBAT, iWAT and eWAT in male UTXKO and fl/Y mice fed HFD (n=3/group). (E) Energy expenditure in male UTXKO and fl/Y mice fed HFD (n=4/group). (F) Thermogenic gene expression in iBAT measured by quantitative RT-PCR in male UTXKO and fl/Y mice fed HFD (n=6/group). (G) Immunoblotting of UCP1 in iBAT of male UTXKO and fl/Y mice fed HFD (n=4/group/group). (H) Immunohistochemistry (IHC) staining of UCP1 in iBAT of male UTXKO and fl/Y mice fed HFD (n=3/group). (I)-(J) Glucose tolerance test (GTT) (I) and Insulin tolerance test (ITT) (J) in male UTXKO and fl/Y mice fed HFD (n=6/group). All data are expressed as mean ± SEM; *p<0.05 vs. fl/Y by two-tailed unpaired Student’s t-test.

### Utx deficiency in Myf5-expressing brown adipocyte precursor cells does not regulate BAT thermogenesis and energy homeostasis

Recent studies suggest that brown adipocytes and skeletal muscle cells share a common developmental lineage and are derived from precursor cells that express myogenic factor 5 (*Myf5*)(Seale et al., 2007). To investigate whether *Utx* regulates brown adipocyte function at an early developmental stage, we generated mice with *Utx* deletion in *Myf5*-expressing brown adipocyte/myotube precursor cells (MUTXKO) by crossing *Utx*-floxed mice (Welstead et al., 2012) with *Myf5-Cre* mice (Tallquist et al., 2000) (Jackson Laboratory, Stock No. 007893). As expected, MUTXKO exhibited around 70% deletion of *Utx* mRNA levels in iBAT (**Suppl. Fig 4A**). However, unlike UTXKO mice with *Utx* deficiency in mature brown adipocytes (**Fig 1**), male mice with *Utx* deficiency in *Myf5*-expressing brown adipocyte/myotube precursor cells had no differences in body weight, body composition, energy expenditure and locomotor activity compared to their fl/Y littermates when fed a HFD diet (**Suppl. Fig 4B-4E**). In addition, there was no difference in *Ucp1* and *Pgc1α* expression in iBAT between male MUTXKO and fl/Y mice on HFD (**Suppl. Fig 4F**). Thus, our data suggest that the regulation of brown adipocyte function by UTX is dependent on brown adipocyte developmental stage; UTX becomes important in maintaining active brown adipocyte function possibly at a later developmental stage, after the precursor cells are committed to the brown adipocyte lineage.

### Utx deficiency in brown fat induces myogenesis

To determine the molecular mechanism whereby *Utx* deficiency impaired brown fat thermogenesis and promoted diet-induced obesity, we performed RNA-seq analysis using iBAT from UTXKO and fl/Y mice fed HFD diet for 12 weeks to unbiasedly examine gene expression pattern changes induced by *Utx* deficiency in brown fat. We found that a total of 1308 genes were differentially regulated by UTX deficiency (Log2 fold change ≥0.5 or ≤-0.5); out of which 254 genes were up-regulated, and 1054 genes were down-regulated by brown adipocyte UTX deficiency. As expected, bioinformatics analysis of these differentially expressed genes with an online software (https://github.com/PerocchiLab/ProFAT)(Cheng et al., 2018) predicted an overall gene expression profile of reduced BAT characteristics in *Utx*-deficient iBAT, with a reciprocal increase in gene expression profile resembling that of WAT (**Fig 2A**). This was consistent with a down-regulation of brown fat-specific gene expression in *Utx*-deficient iBAT (**Fig 2B)**. Surprisingly, a hierarchical cluster analysis disclosed a significant up-regulation of myogenic marker genes in *Utx*-deficient brown fat (**Fig 2C**). Further analysis with quantitative RT-PCR confirmed that myogenic marker gene expression was significantly up-regulated in iBAT of chow-Fed (**Fig 2D**), HFD-fed (**Fig 2E**) or cold-challenged UTXKO mice (**Suppl. Fig 5A**), including skeletal muscle myosin heavy polypeptide 1 (*Myh1*), skeletal muscle myosin heavy polypeptide 4 (*Myh4*), muscle creatine kinase (*Ckm*), sarco/endoplasmic reticulum Ca^2+^-ATPase isoform 1/ATPase, Ca^2+^ transporting, cardiac muscle, fast twitch 1 (*Serca1/Atp2a1*), skeletal muscle α1 actin (*Acta1*), myogenic differentiation 1 (*Myod1)* and myogenin (*Myog*). This was consistent with IHC analysis showing up-regulation of skeletal muscle marker myosin heavy chain (MyHC) in iBAT of UTXKO mice (**Fig 2F**). In addition, primary brown adipocytes isolated from *Utx*-deficient iBAT exhibited reduced oxygen consumption rate (OCR) as measured by Seahorse analyzer (**Fig 2G**), indicating impaired mitochondrial function. Thus, our data revealed an intriguing BAT-to-myocyte remodeling process in BAT of UTXKO mice, which may lead to impaired brown adipocyte mitochondrial function and thermogenesis, thereby contributing to reduced energy expenditure and obesity in UTXKO mice.

**Figure 2.**
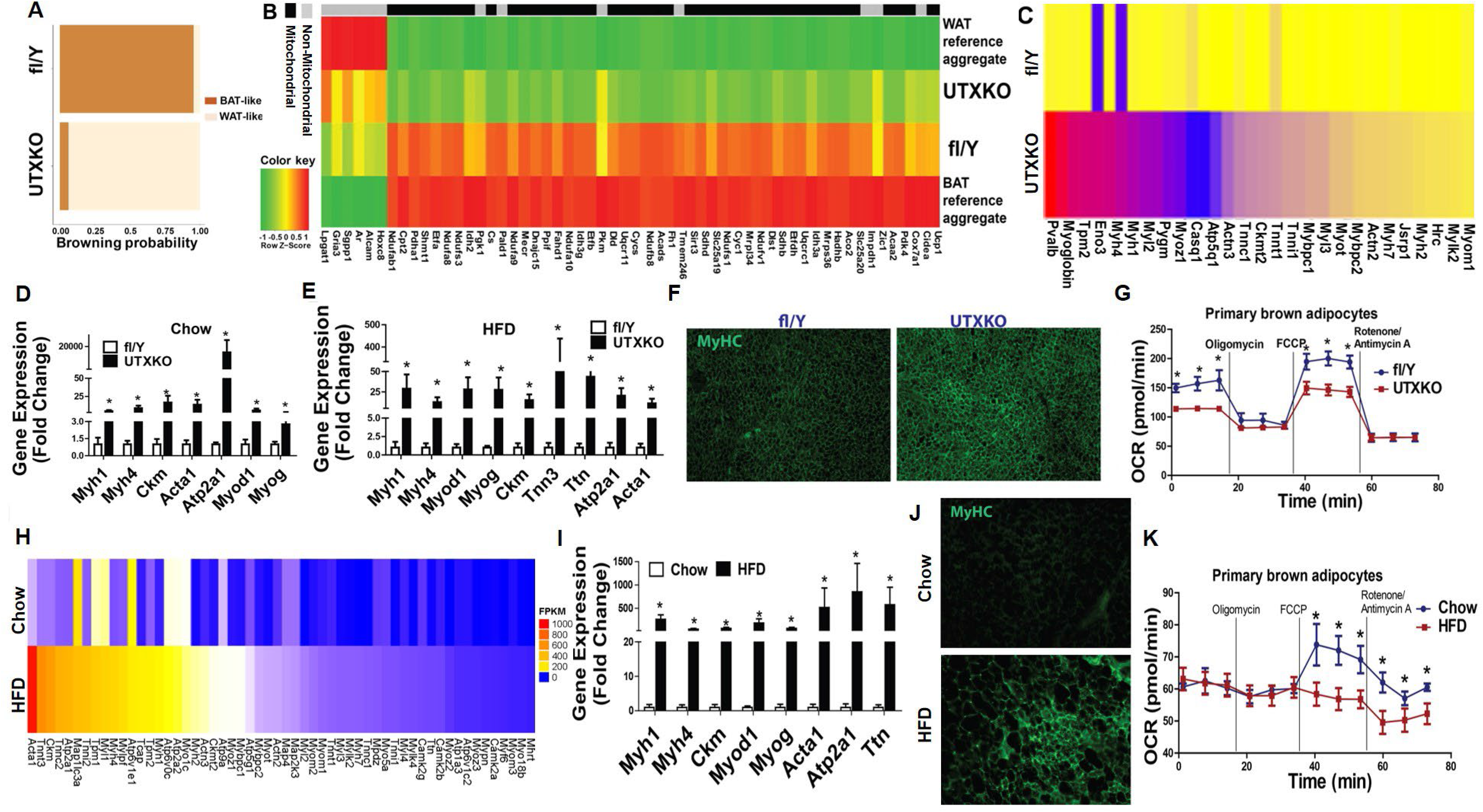
*Utx* deficiency in brown adipocytes induces myogenesis. (A) Bioinformatic modeling of BAT-like or WAT-like gene expression profiles using RNA-seq data from iBAT of male UTXKO and fl/Y mice fed HDF for 12 weeks using an online software (https://github.com/PerocchiLab/ProFAT). (B) RNAseq analysis of BAT-specific gene expression in iBAT of male UTXKO and fl/Y mice fed HDF for 12 weeks using an online software (https://github.com/PerocchiLab/ProFAT). The WAT reference aggregate and BAT reference aggregate were derived from the online software. (C) Heatmap of myogenic marker gene expression in iBAT of male UTXKO and fl/Y mice fed HDF for 12 weeks. (D)-(E) Quantitative RT-PCR analysis of myogenic marker gene expression in iBAT of chow-Fed (D) and HFD-fed (E) UTXKO and fl/Y mice (n=6/group). (F) Immunohistochemistry (IHC) staining of myosin heavy chain (MyHC) in iBAT of UTXKO and fl/Y mice (n=3/group). (G) Oxygen consumption rate (OCR) in primary brown adipocytes isolated from iBAT of male UTXKO and fl/Y mice measured by a Seahorse XF 96 Extracellular Flux Analyzer (n=10/group). (H) Heatmap of myogenic marker gene expression from iBAT of wild type mice fed chow or HFD. (I) Quantitative RT-PCR analysis of myogenic marker gene expression in iBAT of chow- or HFD-fed wild type C57BL/6J mice (n=6/group). (J) IHC staining of MyHC in iBAT of chow- or HFD-fed wild type C57BL/6J mice (n=3/group). (K) OCR of primary brown adipocytes isolated from chow- or HFD-fed wild type C57BL/6J mice (n=10/group). All data are expressed as mean ± SEM; *p<0.05 vs. fl/Y or chow.

Cold and diet are two primary triggers inducing brown fat thermogenesis (Cannon and Nedergaard, 2004, 2011; Feldmann et al., 2009; Lowell and Spiegelman, 2000; von Essen et al., 2017). Diet is a potent stimulator of energy expenditure, a phenomenon referred to as diet-induced thermogenesis that requires both brown fat and UCP1 (Cannon and Nedergaard, 2004; Feldmann et al., 2009; Lowell and Spiegelman, 2000; von Essen et al., 2017). While cold-induced thermogenesis defends body temperature of animals against a cold environment, diet-induced thermogenesis slows down weight gain induced by a short-term overfeeding (Cannon and Nedergaard, 2004; Lowell and Spiegelman, 2000). We therefore explored whether brown fat myogenic remodeling could be modulated during cold- or diet-induced thermogenesis.

Interestingly, cold exposure, a well-known condition that boosts BAT thermogenic function, profoundly suppressed myogenic marker gene expression in iBAT (**Suppl. Fig 5B**). In addition, we also performed a time course study with chow- and HFD-fed animals to further investigate whether such brown adipocyte-myocyte remodeling process also occurred during the course of diet-induced obesity. As shown in **Suppl. Fig 6**, HFD feeding induced *Ucp1* expression in iBAT starting from week 1 and lasted up to 12 weeks when compared to chow-fed animals, suggesting increased diet-induced thermogenesis (**Suppl. Fig 6A**). However, the induction of *Ucp1* by HFD gradually disappeared such that *Ucp1* expression was no longer different between chow and HFD-fed animals after 24 weeks on HFD, suggesting that diet-induced thermogenesis waned after prolonged HFD feeding (**Suppl. Fig 6A**). In contrast, HFD feeding suppressed the expression of myogenic marker genes including *Myod1*, *Myog* and *Myh1* at early stage of HFD feeding from week 1-12, while stimulating the expression of these myogenic marker genes after prolonged HFD feeding (**Suppl. Fig 6B-D**). Further analysis revealed a reciprocal expression pattern of *Ucp1* and myogenic marker genes in iBAT of HFD-fed mice (**Suppl. Fig 6E)**, showing a gradual decline of *Ucp1* expression and a reciprocal increase of myogenic marker gene expression following HFD feeding (**Suppl. Fig 6E**). Indeed, the expression levels of myogenic marker genes were negatively correlated with that of *Ucp1* in iBAT when analyzed from both chow- and HFD-fed mice (**Suppl. Fig 6F**).

To further confirm our findings, we performed RNA-seq analysis in iBAT of wild type mice with chow or HFD feeding for 24 weeks. As expected, prolonged HFD feeding significantly induced myogenic marker gene expression in iBAT of HFD-fed mice compared to that of chow-fed mice (**Fig 2H**). This was further confirmed by quantitative RT-PCR analysis showing up-regulation of myogenic marker gene expression in iBAT of HFD-fed mice, including *Myh1*, *Myh4*, *Ckm*, *Myod1*, *Myog*, *Acta1*, *Atp2a1* and Titin (*Ttn*) (**Fig 2I**). In addition, IHC staining showed a significant upregulation of the skeletal muscle marker MyHC in iBAT of HFD-fed mice compared to that of chow-fed mice (**Fig 2J**), further validating myogenic remodeling of iBAT under prolonged HFD feeding. Importantly, primary brown adipocytes isolated from iBAT of 24-week HFD-fed mice exhibited reduced oxygen consumption rate (OCR) measured by Seahorse analyzer (**Fig 2K**). In consistence, while acute HFD feeding significantly increased oxygen consumption in mice (**Suppl. Fig 7A**), prolonged HFD feeding in mice resulted in a gradual reduction of oxygen consumption compared to that of chow-fed mice (**Suppl. Fig 7B-D**).

Thus, our data revealed a reciprocal BAT-myocyte remodeling process in BAT that is modulated by both cold and diet challenge. The induction of this process by HFD impairs BAT mitochondria function and thermogenesis, which may contribute to reduced energy expenditure and the development of obesity in HFD-fed animals. The fact that both HFD and brown adipocyte-specific *Utx* deletion induce a similar BAT-to-myocyte remodeling indicate that *Utx*-regulated epigenetic modification may be involved in this process.

### DNMT1 deficiency in brown fat induces myogenesis

The interaction of epigenetic mechanisms, including histone methylation and DNA methylation, results in organization of the chromatin structure on different hierarchal levels, which coordinately regulates gene expression (Backdahl et al., 2009; Bannister and Kouzarides, 2011; Maunakea et al., 2010). DNA methylation is catalyzed by DNA methyltransferases (DNMTs). While *de novo* DNA methylation is generally thought to be mediated by DNMT3A and DNMT3B, DNMT1 is believed to function as a maintenance enzyme to maintain DNA methylation status through mitosis using hemi-methylated DNA strands as templates (Backdahl et al., 2009; Bannister and Kouzarides, 2011; Maunakea et al., 2010). However, recent data also suggest that DNMT1 may coordinate with DNMT3A and DNMT3B to regulate *de novo* DNA methylation (Fatemi et al., 2002; Feng et al., 2010; Jeltsch and Jurkowska, 2014; Kim et al., 2002; Okano et al., 1999). Thus, to gain further insight into the epigenetic mechanisms that may regulate the BAT-myocyte remodeling process in brown adipocyte, we have generated mice with brown adipocyte-specific deletion of *Dnmt1*, *Dnmt3a* and *Dnmt3b* to study the role of DNA methylation in this process. Interestingly, we found that mice with brown adipocyte-specific deletion of *Dnmt1* or *Dnmt3a* exhibited similar phenotypes to that of UTXKO (see the results described below), whereas mice with brown adipocyte-specific deletion of Dnmt3b displayed a different metabolic phenotype (Xue and Shi, unpublished data). Thus, we have focused on brown adipocyte DNMT1 and DNMT3A in the current study and further explored whether UTX interacts with DNMTs in the regulation of BAT-myocyte remodeling in brown fat.

To generate mice with brown fat-specific deletion of *Dnmt1* (D1KO), we crossed *Dnmt1*-floxed mice (fl/fl) (Jackson-Grusby et al., 2001) with *Ucp1*-*Cre* mice (Kong et al., 2014). We found *Dnmt1* deletion in brown fat resulted in around 50% reduction of *Dnmt1* mRNA expression (**Suppl. Fig 8A**). Interestingly, RNA-seq analysis using iBAT from D1KO and fl/fl mice revealed a significant up-regulation of myogenic gene expression in *Dnmt1*-deficient brown fat (**Fig 3A**), similarly to that observed in iBAT of HFD-fed and UTXKO mice. This was further confirmed by quantitative RT-PCR analysis showing up-regulation of key myogenic gene expression, including *Myod1*, *Myog*, *Myh1*, *Myh4*, *Ckm*, *Serca1/Atp2a1*, *Ttn*, and *Acta1* in iBAT of D1KO mice (**Fig 3B**). More importantly, bioinformatic analysis of RNAseq data from iBAT of HFD-fed, UTXKO and D1KO revealed a group of myogenic marker genes that were similarly up-regulated in all three datasets (**Suppl. Fig 8B**), indicating that both DNMT1 and UTX might be involved in the regulation of BAT-to-myocyte remodeling in iBAT of HFD-fed mice.

**Figure 3.**
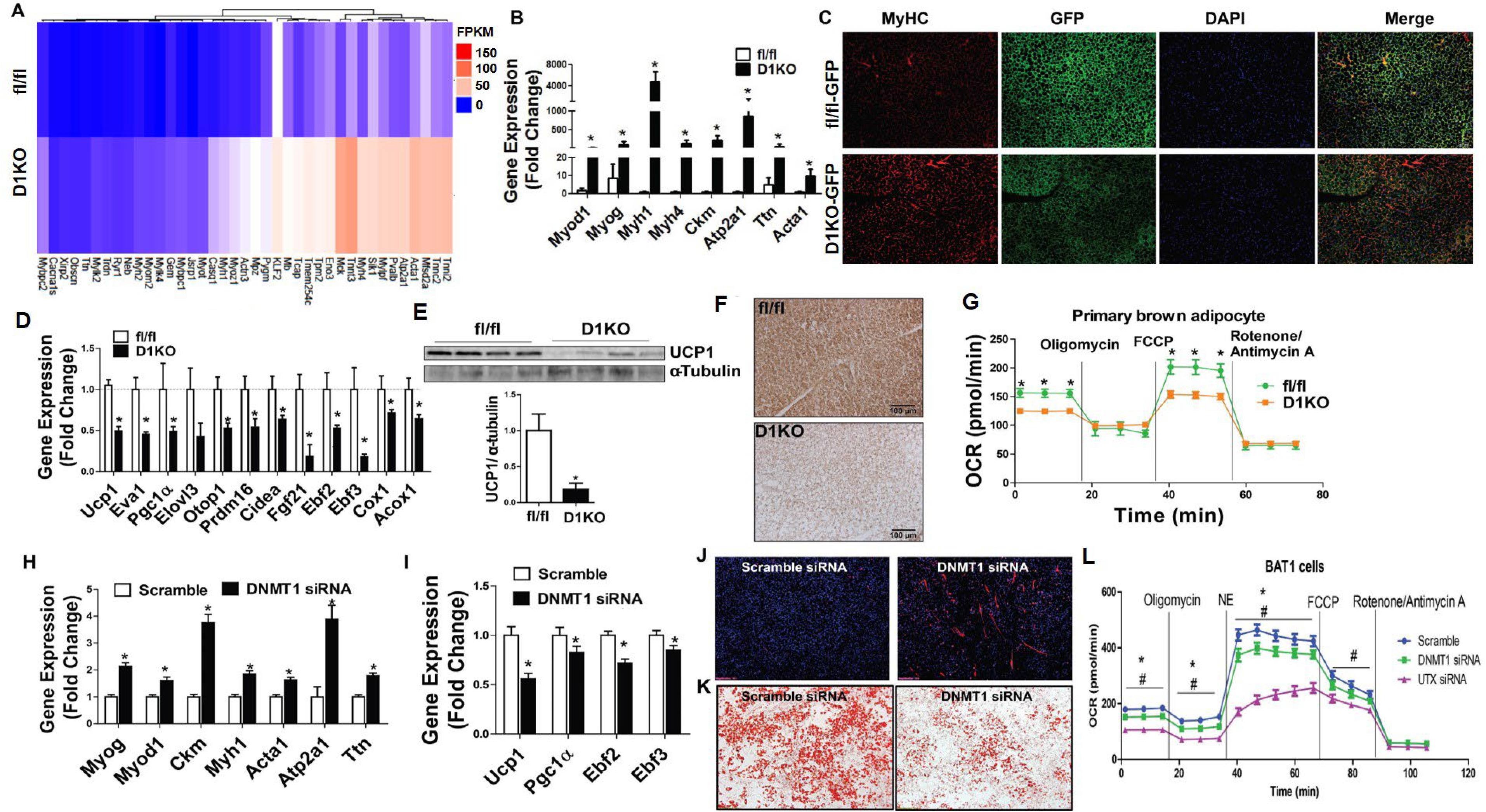
*Dnmt1* deficiency in brown fat induces myogenesis. (A) Heatmap of myogenic gene expression in iBAT from D1KO and fl/fl mice on regular chow diet. (B) Quantitative RT PCR analysis of myogenic gene expression in D1KO and fl/fl mice on chow diet (n=6/group). (C) IHC staining of MyHC in GFP-labeled brown adipocytes in iBAT of D1KO-GFP and fl/fl-GFP mice on chow diet (n=3/group). (D) Quantitative RT-PCR analysis of BAT-specific gene expression in iBAT of D1KO and fl/fl mice on chow diet (n=6/group). (E) Immunoblotting analysis of UCP1 protein levels in iBAT of D1KO and fl/fl mice on chow diet (n=4/group). (F) IHC staining of UCP1 in iBAT of D1KO and fl/fl mice on chow diet (n=3/group). (G) OCR of primary brown adipocytes isolated from iBAT of D1KO or fl/fl mice measured by a Seahorse analyzer (n=10/group). (H)-(I) Quantitative RT-PCR analysis of myogenic gene expression (H) and BAT-specific gene expression (I) in BAT1 brown adipocytes transfected with scramble or *Dnmt1* siRNA (n=10/group). (J)-(K) IHC staining of MyHC (J) and Oil red O staining (K) in BAT1 brown adipocytes transfected with scramble or *Dnmt1* siRNA (n=3/group). (L) OCR in BAT1 brown adipocytes transfected with scramble, *Dnmt1* or *Utx* siRNA measured by a Seahorse analyzer (n=10/group). For (H)-(L), BAT1 cells were differentiated into brown adipocytes as described under Methods and scramble and targeting siRNAs were transfected into day 4 differentiated BAT1 cells using Amaxa Nucleofector II Electroporator with an Amaxa cell line nucleofector kit L according to the manufacturer’s instructions (Lonza). Cells were harvested 2 days after for further analysis. All data are expressed as mean ± SEM. *p<0.05 vs. fl/fl or Scramble siRNA by two-tailed unpaired Student’s t-test. For L, *p<0.05 scramble vs. DNMT1 siRNA, #p<0.05 scramble vs. UTX siRNA. by ANOVA followed by Bonferroni post-hoc analysis.

We also generated mice with brown adipocytes-specific expression of GFP by triple-crossing *Dnmt1*-floxed mice, *Rosa-Gfp* mice (Srinivas et al., 2001) and *Ucp1*-Cre mice (D1KO-GFP, or *Ucp1-Cre::Dnmt1^fl/fl^: Rosa-Gfp^fl/fl^*). IHC staining indicated that *Dnmt1* deficiency markedly induced skeletal muscle marker MyHC in GFP-labeled brown adipocytes in D1KO-GFP mice (**Fig 3C**). In addition, there was a reciprocal down-regulation of brown fat thermogenic gene expression in iBAT from D1KO mice (**Fig 3D**), which was associated with decreased UCP1 protein content and UCP1 IHC staining in iBAT of D1KO mice (**Fig 3E** and **3F**). Similar to that of *Utx*-deficient iBAT, the BAT-to-myocyte remodeling in *Dnmt1*-deficient brown adipocytes resulted in reduced oxygen consumption rate (**Fig 3G**), indicative of impaired mitochondrial function. On the other hand, bioinformatic analysis of RNA-seq data with an online software (Cheng et al., 2018) predicted that iBAT from D1KO mice exhibited gene expression profiles characteristic of that of white fat (**Suppl. Fig 8C**), with less expression of BAT- and mitochondria-enriched genes (**Suppl. Fig 8D**).

To further study whether *Dnmt1* regulated BAT-to-myocyte remodeling via a cell autonomous manner, we knocked down *Dnmt1* in 4-day differentiated BAT1 brown preadipocytes via siRNA approach. As expected, knockdown of *Dnmt1* in BAT1 cells significantly promoted myogenic gene expression, including *Myog*, *Myod1*, *Ckm*, *Myh1*, *Acta1*, *Atp2a1* and *Ttn* (**Fig 3H**) while simultaneously down-regulating BAT-specific gene expression, including *Ucp1*, *Pgc1α*, *Ebf2* and *Ebf3* (**Fig 3I**). Moreover, knockdown of *Dnmt1* enhanced myogenesis as evidenced by increased MyHC staining (**Fig 3J**), while simultaneously reducing brown adipogenesis as evidenced by much less neutral lipid accumulation measured by Oil red O staining (**Fig 3K**). In consistence, Seahorse analysis revealed a decreased OCR in *Dnmt1* knockdown BAT1 brown adipocytes and similarly but to a greater extent, in *Utx* knockdown BAT1 brown adipocytes (**Fig 3L**).

### Mice with Dnmt1 deficiency in brown fat exhibit increased body weight and adiposity on chow diet and are susceptible to DIO when fed a HFD

To determine the role of brown adipocyte DNMT1 in the regulation of body weight and energy metabolism in animal models, we conducted metabolic characterization in D1KO mice. As shown in **Fig 4A**, female D1KO mice gained more weight even on a regular chow diet compared to their fl/fl littermate control mice. In consistence, D1KO mice exhibited a significant increase in fat mass and a significant decrease in lean mass (**Fig 4B**). Individual fat pad mass including iWAT, gonadal WAT (gWAT) and rWAT was also increased in D1KO compared to that of fl/fl mice (**Fig 4C**). In addition, histological examination revealed larger adipocyte size in iBAT, iWAT and gWAT of female D1KO mice (**Fig 4D**). Using PhenoMaster metabolic cage systems, we found that female D1KO mice displayed lower energy expenditure with decreased oxygen consumption (**Fig 4E** and **Suppl. Fig 9A**), while there was no change in RER, locomotor activity, and a tendency of lower food intake in D1KO (**Suppl. Fig 9B**-**9D**). Female D1KO mice also displayed insulin resistance on the regular chow diet, as shown by increased fasting and fed insulin levels (**Fig 4F**) and impaired glucose tolerance and insulin sensitivity in GTT and ITT tests (**Fig 4G** and **4H**).

**Figure 4.**
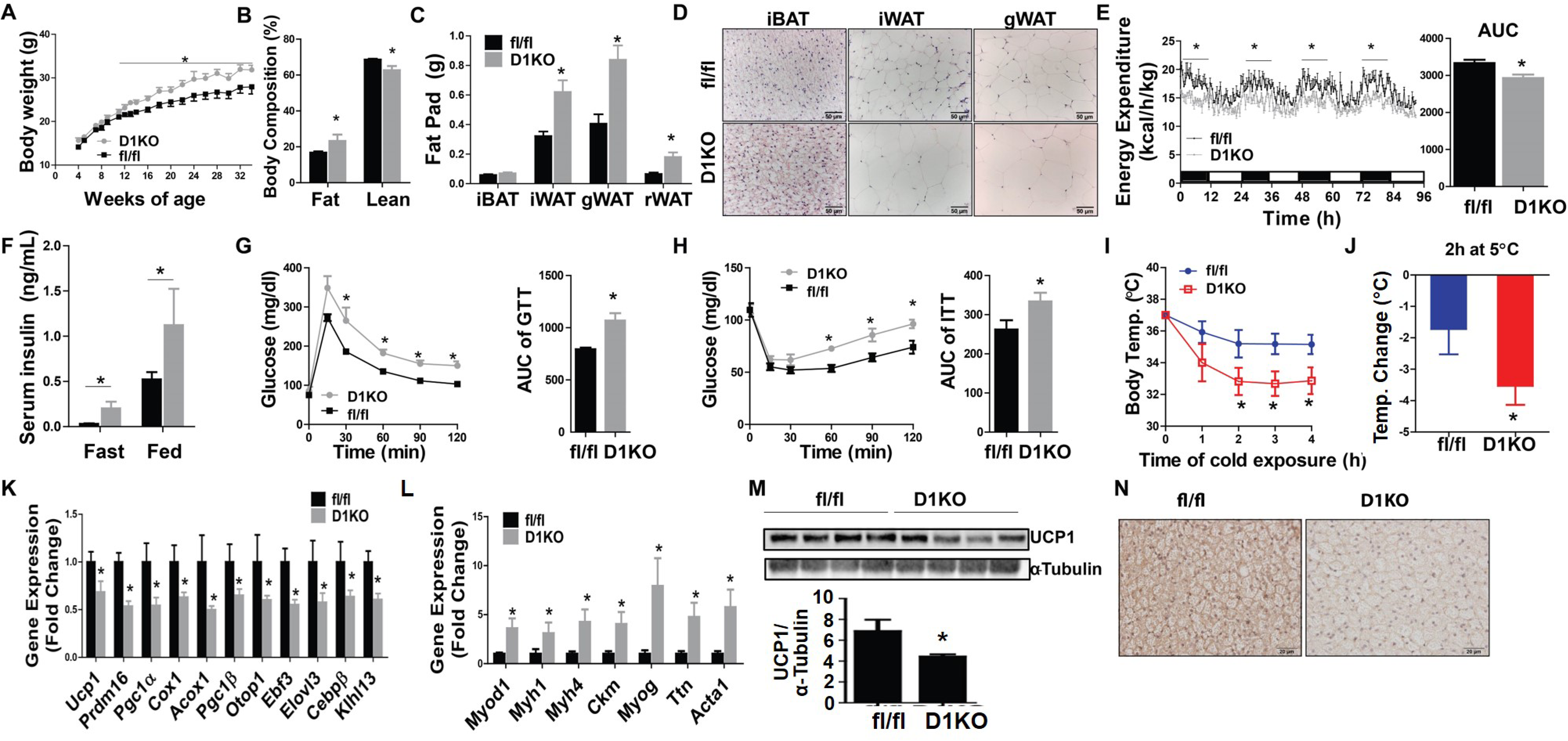
DNMT1 deficiency in brown fat impairs brown fat function. D1KO and fl/fl mice were weaned onto regular chow diet and various metabolic phenotypes were studied. (A)-(C) Body weight growth curve (A), Body composition (B), and Fat pad weight (C) in female D1KO and fl/fl mice on chow diet (n=6-8/group). (D) H&E staining of iWAT and iBAT of female D1KO and flf/fl mice on chow diet (n=4/group). (E) Energy expenditure in female D1KO and fl/fl mice on chow diet (n=4/group). (F)-(H) Fasting and fed serum insulin levels (F), GTT (G), and ITT (H) of female D1KO and fl/fl mice on chow diet (n=6-8/group). (I) Body temperature in male 2-month-old D1KO and fl/fl mice during acute 5°C cold exposure (n=6/group). (J) Body temperature change in male 2-month-old D1KO and fl/fl mice after 2 hours of cold exposure (n=6/group). (K)-(L) Quantitative RT-PCR analysis of BAT-specific gene expression (K), and myogenic gene expression (L) in iBAT of male 2-month-old D1KO and fl/fl mice after an acute 4-hour 5°C cold exposure (n=6/group). (M) Immunoblotting of UCP1 protein in iBAT of male 2-month-old D1KO and fl/fl mice after a 7-day 5°C cold exposure (n=4/group). (N) UCP1 IHC staining in iBAT from male 2-month-old D1KO and fl/fl mice after a chronic 5°C 7-day cold exposure (n=3/group). All data are expressed as mean ± SEM. *p<0.05 by two-tailed unpaired Student’s t-test.

Similar results were also observed in male D1KO mice. When fed a regular chow diet, male D1KO mice also exhibited increased body weight and adiposity compared to their fl/fl littermate control mice (**Suppl. Fig 10A** and **10B**). They also displayed glucose intolerance and insulin resistance in GTT and ITT tests compared to fl/fl mice (**Suppl. Fig 10C** and **10D**).

To determine the role of brown adipocyte DNMT1 in diet-induced obesity, we challenged male and female D1KO mice and their littermate control fl/fl mice with HFD for up to 24 weeks. As expected, HFD-fed female D1KO mice gained more weight compared to fl/fl mice (**Suppl.** F**ig 11A**). Body composition analysis revealed a significant increase in fat mass and a significant decrease in lean mass (**Suppl. Fig 11B**), which was associated with increased individual fat pad mass in iWAT, gWAT and iBAT (**Suppl. Fig 11C**). Similar to chow-fed animals, female D1KO mice on HFD also exhibited reduced energy expenditure with decreased oxygen consumption (**Suppl. Fig 11D-11E**) without changes in RER, locomotor activity and food intake (**Suppl. Fig 11F-11H**). With the increased adiposity, D1KO mice had increased fed insulin levels (**Suppl. Fig 11I**) and displayed glucose intolerance and insulin resistance as assessed by GTT and ITT (**Suppl. Fig 11J** and **11K**), respectively.

Similarly, male D1KO mice gained more weight compared to their fl/fl littermate control mice upon HFD challenge (**Suppl. Fig 12A**), which was associated with increased fat mass in iWAT (**Suppl. Fig 12B**) and larger adipocytes in iWAT and iBAT (**Suppl. Fig 12C**). Male D1KO mice on HFD also exhibited lower energy expenditure with decreased oxygen consumption (**Suppl. Fig 12D-12E**) without changes in RER and locomotor activity (**Suppl. Fig 12F-12G**). In addition, male D1KO mice had slightly decreased cumulative food intake (**Suppl. Fig 12H**), possibly due to adaptation to their reduced energy expenditure. Further, male D1KO mice exhibited down-regulated thermogenic gene expression in iBAT and iWAT (**Suppl. Fig 12I** and **12J**), with decreased UCP1 protein levels in iBAT (**Supp. Fig 12K**). Consistent with their increased adiposity, male D1KO mice displayed glucose intolerance and insulin resistance as assessed by GTT and ITT (**Suppl. Fig 12L** and **12M**). Thus, our data consistently indicate that DNMT1 deficiency in brown fat promotes obesity and impairs insulin sensitivity in mice.

### DNMT1 deficiency in brown fat impairs cold-induced thermogenesis

To study the role of brown fat DNMT1 in cold-induced thermogenesis, we challenged D1KO and fl/fl mice for either an acute (4-hour) or a chronic (7-day) cold exposure at 5°C. Interestingly, D1KO mice exhibited lower body temperature during acute cold exposure (**Fig 4I-4J**), with bigger temperature reduction than control fl/fl mice within the first 2 hours of cold exposure (**Fig 4J**), suggesting that the DNMT1-deficient mice are cold sensitive. During the acute cold exposure, Dnmt1 deficiency in iBAT resulted in reduced BAT-specific gene expression, with simultaneously up-regulated myogenic marker gene expression (**Fig 4K-4L**). Similarly, during the chronic cold exposure, DNMT1 deficiency in iBAT also resulted in reduced expression of BAT-specific genes, while reciprocally increasing the expression of myogenic marker genes in iBAT of D1KO mice (**Suppl. Fig 13A-13B**). In consistence with gene expression profiles, iBAT from D1KO mice also exhibited decreased UCP1 protein level (**Fig 4M**), larger brown adipocytes as measured by H&E staining (**Suppl. Fig 13C**), and less UCP1 IHC staining in response to the chronic cold exposure (**Fig 4N**). These data suggest that DNMT1 deficiency in brown fat impairs cold-induced thermogenesis.

### Mice with brown adipocyte-specific Dnmt3a deficiency displayed similar metabolic phenotypes to that of UTXKO and D1KO

We also generated mice with brown fat-specific deletion of *Dnmt3a*, a DNA methyltransferase that is responsible for *de novo* DNA methylation (D3aKO), by crossing *Dnmt3a*-floxed mice (Kaneda et al., 2004) with *Ucp1-Cre* mice (Kong et al., 2014), and found that D3aKO mice had similar metabolic phenotypes to that of D1KO. For instance, female D3aKO mice gained more weight even on regular chow diet compared to their fl/fl littermate control mice (**Suppl. Fig 14A**) even with less food intake (**Suppl. Fig 14B**). In consistence, D3aKO mice exhibited larger fat pads and liver compared to that of fl/fl mice (**Suppl. Fig 14C**). Using PhenoMaster metabolic cage systems, we found that D3aKO mice displayed lower energy expenditure (**Suppl. Fig 14D**) with decreased oxygen consumption (**Suppl. Fig 14E**) with largely no changes in locomotor activity (**Suppl. Fig 14F**). Further GTTs and ITTs confirmed that D1KO mice developed glucose intolerance and insulin resistance (**Suppl. Fig 14G** and **14H**).

### Myod1 mediates the effect of DNMT1 deficiency on brown fat myogenesis

To further determine potential myogenic transcriptional factors whose up-regulations are direct targets by promoter demethylation due to *Dnmt1* deficiency, we performed a DNA methylation profiling experiment using Reduced Representation Bisulfite Sequencing (RRBS) approach (Bock et al., 2010; Meissner et al., 2005; Smith et al., 2009). Since our data suggest that iBAT from female D1KO mice exhibits prominent BAT-to-myocyte remodeling compared to their fl/fl littermates even on a regular chow diet, we used DNA samples from iBAT of chow-fed female D1KO and fl/fl mice for the RRBS analysis. We found there are up to 242 Differentially Methylated Regions (DMRs) in iBAT of D1KO vs fl/fl mice. Within these DMRs, around 202 were located within genes, with the rest associated with gene promoters. Notably, our RRBS profiling indicated that the methylation rate at the proximate promoter and 5’-region of *Myod1* gene was significantly decreased in iBAT of D1KO mice compared to that of fl/fl mice (**Fig 5A**), whereas *Myd1* express was reciprocally up-regulated in iBAT from UTXKO, HFD-fed and D1KO mice (**Fig 2E, 2I** and **3B**). Interestingly the proximal promoter and 5’ region of *Myod1* are enriched with CpG sites (**Suppl. Fig 15A**); alterations of DNA methylation on these CpG sites could potentially alter *Myod1* expression.

**Figure 5.**
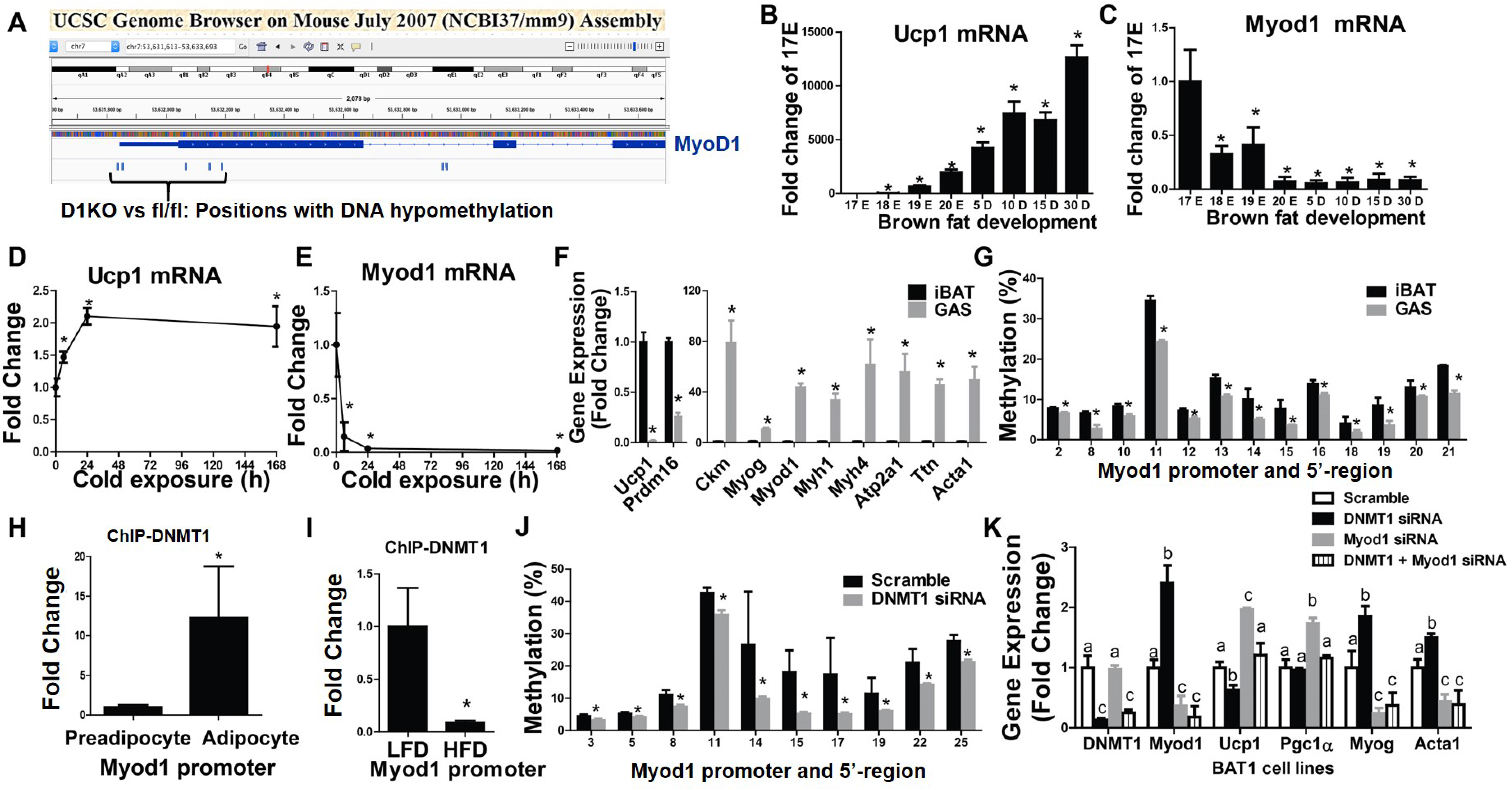
MyoD1 mediates the effect of DNMT1 deficiency on brown fat myogenesis. (A) RRBS profiling of DNA methylation level at MyoD1 promoter in iBAT of D1KO and fl/fl mice. (B)-(C) *Ucp1* (B) and *Myod1* (C) expression in iBAT of mice during late embryonic and postnatal development (n=4/group). (D)-(E) *Ucp1* (D) and *Myod1* (E) expression in iBAT of mice during cold exposure (n=4/group). (F) *Ucp1*, *Prdm16* and myogenic marker gene expression in iBAT and gastrocnemius (GAS) muscle (n=4/group). (G) Pyrosequencing analysis of DNA methylation level at *Myod1* promoter in iBAT and GAS muscle (n=4/group). (H) ChIP assay of DNMT1 binding to *Myod1* promoter in undifferentiated BAT1 preadipocytes and differentiated BAT1 brown adipocytes (n=3/group). (I) ChIP assay of DNMT1 binding to the *Myod1* promoter in iBAT from HFD- or LFD-fed mice (n=4/group). (J) Pyrosequencing analysis of DNA methylation levels at *Myod1* promoter in BAT1 brown adipocytes transfected with scramble or *Dnmt1* siRNA (n=6/group). (K) Quantitative RT-PCR analysis of myogenic marker gene and BAT gene expression in BAT1 brown adipocytes transfected with scramble, *Dnmt1*, *Myod1*, or *Dnmt1*+*Myod1* siRNA (n=4/group). Different lowercase letters represent significant differences at p<0.05. For (J) and (K), BAT1 cells were differentiated into brown adipocytes as described under Methods and scramble and targeting siRNAs were transfected into day 4 differentiated BAT1 cells using Amaxa Nucleofector II Electroporator with an Amaxa cell line nucleofector kit L. Cells were harvested 2 days after for pyrosequencing (J) or gene expression (K) analysis. All data are expressed as mean ± SEM. *p<0.05 by Student’s t-test or ANOVA with Bonferroni post-hoc analysis.

Prior lineage tracing studies show that brown fat and skeletal muscle indeed share the same developmental origins (Seale et al., 2008). To determine the role of *Myod1* in the determination of brown fat and myocyte lineage, we interrogated the relationship between the myogenic driver *Myod1* and the BAT marker *Ucp1* by measuring the expression pattern of these two genes during brown fat development. *Ucp1* expression in iBAT began to increase at embryonic day 17 (E17), and continued to rise postnatally (**Fig 5B**), similar to previous reports (Barnd et al., 1970; Skala et al., 1970; Xue et al., 2007). In contrast, *Myod1* expression was at the highest level in iBAT at E17, which then sharply declined afterwards and stayed at low levels postnatally (**Fig 5C**). These data suggest that *Myod1* and *Ucp1* expression is inversely correlated and may be mutually exclusive. Moreover, reciprocal regulation of *Myod1* and *Ucp1* expression was also evident in cold exposure such that cold exposure markedly stimulated *Ucp1* expression (**Fig 5D**) within 24 hours while simultaneously down-regulating *Myod1* expression (**Fig 5E**). The enrichment of myogenic genes such as *Myod1* along with other markers is a molecular signature of the skeletal muscle, which distinguishes gastrocnemius muscle from iBAT (**Fig 5F**). Interestingly, the high expression of *Myod1* in skeletal muscle was associated with decreased methylation rate at a number of CpG sites at the *Myod1* promoter and 5’-region as measured by pyrosequencing analysis (**Fig 5G**). Hypomethylation at the *Myod1* promoter and 5’-region may contribute to higher *Myod1* gene expression in skeletal muscle, which may be important in maintaining myogenic signature in skeletal muscle.

We then further determined the role of DNMT1 in *Myod1* promoter methylation. Our ChIP assay revealed a significantly higher DNMT1 binding to the *Myod1* promoter in differentiated brown adipocytes BAT1 compared to undifferentiated BAT1 preadipocytes (**Fig 5H**), indicating that enhanced methylation at *Myod1* promoter due to DNMT1 binding may decrease *Myod1* expression in mature brown adipocytes. Further, down-regulated binding of DNMT1 to *Myod1* promoter was also observed in iBAT from HFD fed mice compared to LFD-fed mice (**Fig 5I**). Decreased DNMT1 binding to *Myod1* promoter may increase its gene expression, resulting in the induction of myogenic program seen in the iBAT from HFD fed mice (**Fig 2H-K,** and **Suppl. Fig 6**). To determine whether the effect of DNMT1 on *Myod1* promoter methylation was via a cell autonomous action, we knocked down *Dnmt1* in BAT1 cells. We found that DNMT1 knockdown in BAT1 brown adipocytes resulted in reduced methylation at a number of CpG sites at *Myod*1 promoter (**Fig 5J**).

To confirm our findings that *Myod1* serves as an epigenetic target for *Dnmt1* and mediates *Dnmt1* deletion-induced BAT-to-myocyte remodeling in brown adipocytes, we knocked down *Dnmt1* and *Myod1* individually or in combination in BAT1 brown adipocytes and measured BAT- and myogenic-specific gene expression. As expected, BAT1 brown adipocytes transfected with *Dnmt1* and/or *Myod1* siRNA had significantly reduced *Dnmt1* and/or *Myod1* expression, respectively, confirming the knockdown efficiency (**Fig 5K**). Consistent with the decreased methylation at *Myod1* promoter by *Dnmt1* knockdown (**Fig 5J**), *Dnmt1* knockdown in BAT1 brown adipocytes significantly up-regulated *Myod1* expression, which was abolished by *Myod1* knockdown (**Fig 5K**). Interestingly, *Dnmt1* knockdown in BAT1 brown adipocytes down-regulated BAT-specific *Ucp1* and *Pgc1α* expression while up-regulating myogenic gene expression, including *Myog* and *Acta1* (**Fig 5K**). This was completely reversed by *Myod1* knockdown (**Fig 5K**). Thus, our *in vitro* data confirm our findings that DNMT1 determines BAT-myocyte remodeling via *Myod1* promoter methylation in brown adipocytes.

### Specifically reducing DNA methylation at Myod1 promoter induces myocyte-like brown adipocytes

We next utilized a targeted demethylation approach to determine whether specifically targeting CpG sites and reducing their methylation status at *Myod1* promoter could recapitulate *Dnmt1* deletion-induced BAT-to-myocyte remodeling in brown adipocytes. To achieve site-specific demethylation at *Myod1* promoter, we employed a modified endonuclease dead version of CRISPR associated protein 9 (dCas9) fused with the catalytic domain (CD) of the enzyme involved in DNA demethylation, tet methylcytosine dioxygenase 1 (TET1) (dCas9-TET1CD) (Amabile et al., 2016; Liu et al., 2016; Morita et al., 2016; Vojta et al., 2016; Xu et al., 2016) (**Suppl. Fig 15B**). We transfected BAT1 brown adipocytes with plasmids expressing dCas-TET1CD along with plasmids expressing either scramble non-targeting guide RNA (scramble-gRNA-mCherry) or gRNA targeting CpG sites at *Myod1* promoter (*Myod1*-gRNA-mCherry), and found that DNA methylation rate of a number of CpG sites at *Myod1* promoter was significantly reduced (**Fig 6A**). The down-regulated DNA methylation at *Myod1* promoter was associated with up-regulation of *Myod1* mRNA expression and reciprocal down-regulation of thermogenic gene expression such as *Ucp1*, *Pgc1α*, *Ebf2* and *Prdm16* (**Fig 6B**), suggesting that we have successfully established an approach to specifically target DNA methylation at *Myod1* promoter.

**Figure 6.**
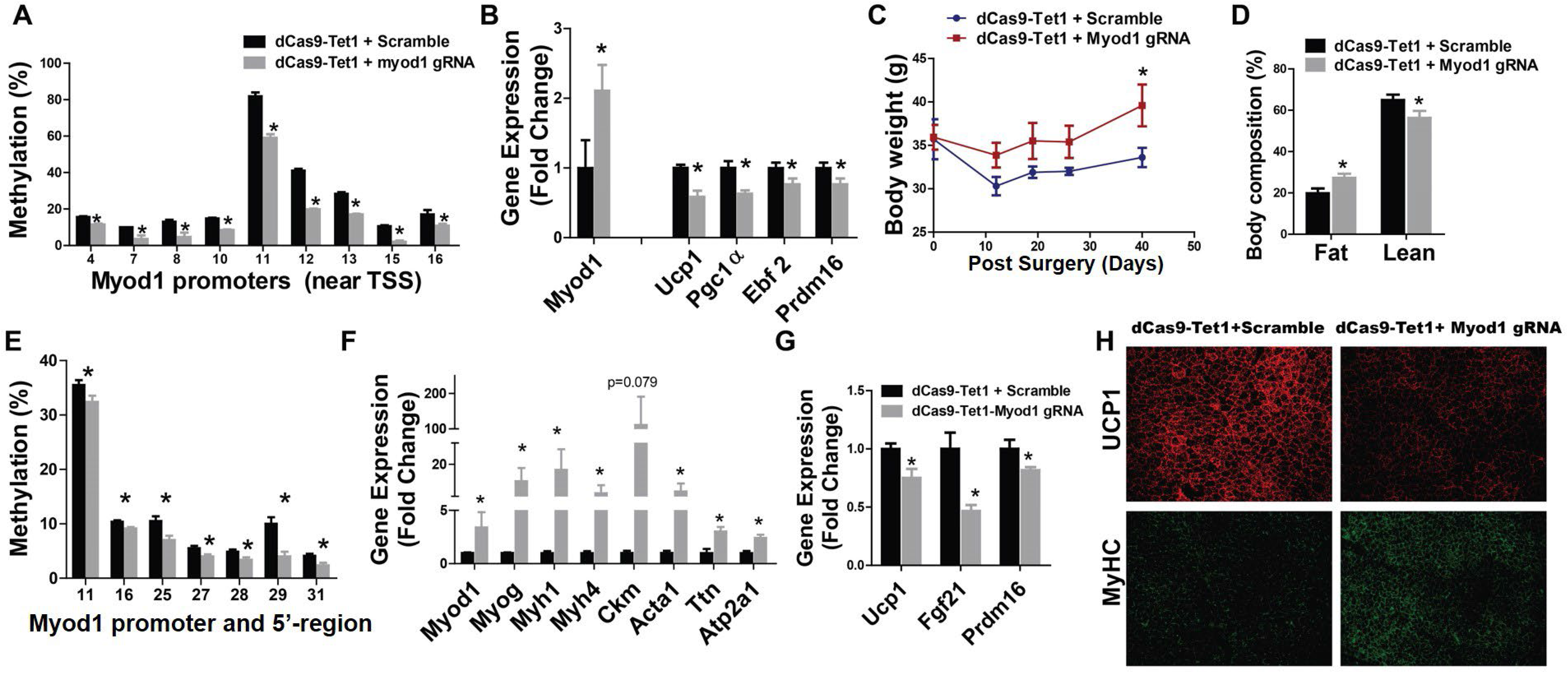
Specifically reducing DNA methylation at *Myod1* promoter in iBAT of mice induces BAT-to-myocyte switch. (A) Pyrosequencing analysis of DNA methylation at *Myod1* promoter in BAT1 adipocytes transfected with lentiviral vectors expressing dCas9-TET1CD along with lentiviral vectors expressing either *Myod1*-targeting gRNA or scramble non-targeting gRNA (n=4/group). (B) Quantitative PCR analysis of Myod1 and BAT-specific gene expression in BAT1 adipocytes transfected with lentiviral vectors expressing dCas9-TET1CD along with lentiviral vectors expressing either *Myod1*-targeting gRNA or scramble non-targeting gRNA (n=4/group). For (A)-(B), 4-day differentiated BAT brown adipocytes were transfected with lentiviral vectors FUW-dCas9-TET1CD along with lentiviral vectors pgRNA-mCherry encoding either scramble-gRNA or Myod1-targeting gRNA using Amaxa Nucleofector II Electroporator with an Amaxa cell line nucleofector kit L. Cells were harvested 2 days after for pyrosequencing (A) or gene expression (B) analysis. (C)-(D) Body weight (C) and Body composition (D) in mice with iBAT injection of lentiviruses expressing dCas9-TET1CD plus lentiviruses expressing either targeting Myod1-gRNA-mCherry or non-targeting scramble-gRNA-mCherry (n=4/group). (E)-(G) Methylation levels at *Myod1* promoter (E), Myogenic marker gene expression (F), and BAT-specific gene expression (G) in iBAT of mice with iBAT injection of lentiviruses expressing dCas9-TET1CD plus lentiviruses expressing either targeting Myod1-gRNA-mCherry or non-targeting scramble-gRNA-mCherry (n=6/group). (H) IHC staining of UCP1 (upper panel) and MyHC (lower panel) in iBAT of mice with iBAT injection of lentiviruses expressing dCas9-TET1CD plus lentiviruses expressing either targeting Myod1-gRNA-mCherry or non-targeting scramble-gRNA-mCherry (n=3/group). For (C)-(H), 3-month-old chow-fed male C57BL/6J mice were bilaterally injected with lentiviruses expressing dCas9-TET1CD plus lentiviruses expressing either targeting Myod1-gRNA-mCherry or non-targeting scramble-gRNA-mCherry into iBAT for up to 2 months. All data are expressed as mean ± SEM. *p<0.05 by Student’s t-test.

To study the role of targeted DNA methylation at *Myod1* promoter in brown fat *in vivo*, we surgically injected lentiviruses expressing dCas9-TET1CD along with lentiviruses expressing either scramble-gRNA-mCherry or *Myod1*-gRNA-mCherry bilaterally into iBAT of male C57BL/6J mice on chow diet. IHC analysis with mCherry and perilipin antibodies confirmed successful infection of the lentiviruses into iBAT brown adipocytes (**Suppl. Fig 16**). The mice receiving lentiviral injection of dCas9-TET1CD and *Myod1*-targeting gRNA exhibited higher body weight (**Fig 6C**) and body fat composition (**Fig 6D**) even on chow diet, thus mimicking the metabolic phenotypes of male and female D1KO mice on chow diet (**Fig 4 and Suppl. Fig 10**). Moreover, lentiviral treatment of dCas9-TET1CD significantly reduced DNA methylation rate of a number of CpG sites at *Myod1* promoter (**Fig 6E**). The decreased methylation at *Myod1* promoter appeared to result in a marked induction of myogenic gene expression (**Fig 6F**), which was associated with a down-regulation of thermogenic gene expression (**Fig 6G**). Further IHC staining revealed a significant upregulation of the skeletal muscle marker MyHC and a reciprocal down-regulation of the thermogenic marker UCP1 (**Fig 6H**). These data indicate that decreased DNA methylation at *Myod1* promoter initiates a BAT-to-myocyte remodeling process in brown fat.

### DNMT1 silences Myod1 expression via interacting with PRDM16

We next determined the molecular mechanism underlying the strikingly similar BAT-to-myocyte remodeling phenotype in BAT of UTXKO and D1KO mice. *Prdm16* and *Myod1* are mutually exclusive in lineage determination of brown adipocytes vs. myocytes (Sanchez-Gurmaches and Guertin, 2014). We found that *Prdm16* expression was down-regulated in iBAT of HFD-fed mice (**Fig 7A**), This may be due to a decreased binding of UTX to *Prdm16* promoter as shown in ChIP assays (**Fig 7B**). Conversely, the level of H3K27me3, which was regulated by UTX (Ge, 2012), was increased at *Prdm16* promoter (**Fig 7C**). To confirm whether *Prdm16* expression is indeed regulated by UTX through modulating of H3K27me3 levels, we knocked down *Utx* in BAT1 brown adipocytes and measured H3K27me3 at *Prdm16* promoter via ChIP assay following stimulation by the β-adrenergic agonist isoproterenol. We found that H3K27me3 level was increased at *Prdm16* promoter in isoproterenol-treated BAT1 brown adipocytes with *Utx* knockdown (**Fig 7D**). Thus, our data suggest that UTX promotes *Prdm16* expression via demethylating H3K27me3 at *Prdm16* promoter.

**Figure 7.**
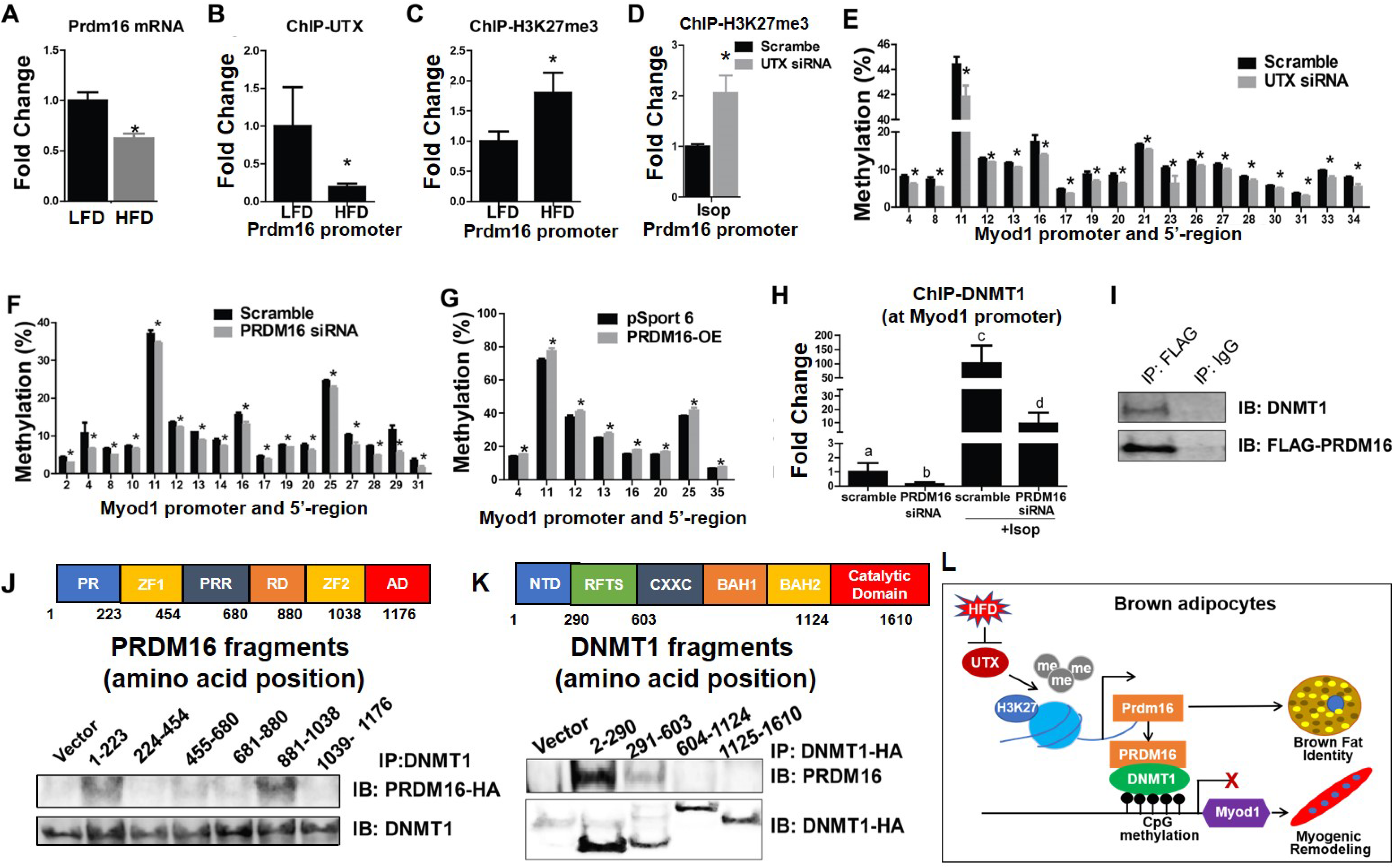
DNMT1 silences *Myod1* expression via interacting with PRDM16. (A) Quantitative RT-PCR analysis of *Prdm16* mRNA in iBAT of LFD- or HFD-fed mice (n=8/Group). (B)-(C) ChIP assay of UTX binding to *Prdm16* promoter (B) and ChIP assay of H3K27me3 levels at *Prdm16* promoter (C) in iBAT of LFD- or HFD-fed mice (n=4/Group). (D) ChIP assay of H3K27me3 levels at *Prdm16* promoter in control or *Utx* knockdown BAT1 brown adipocytes treated with isoproterenol (n=4/Group). (E) Pyrosequencing analysis of DNA methylation at *Myod1* promoter in BAT1 brown adipocytes transfected with scramble or *Utx* siRNA (n=6/Group). (F) Pyrosequencing analysis of DNA methylation at *Myod1* promoter in BAT1 brown adipocytes transfected with scramble or *Prdm16* siRNA (n=4/group). (G) Pyrosequencing analysis of DNA methylation at *Myod1* promoter in BAT1 brown adipocytes transfected with pSport6 or pSport6 encoding *Prdm16* overexpressing plasmids (n=4/group). (H) ChIP assay of DNMT1 binding to *Myod1* promoter in control or *Prdm16* knockdown BAT1 brown adipocytes treated with or without isoproterenol (n=4/group). Different lowercase letters represent significant differences at *p<0.05.* (I) Co-IP of DNMT1 and FLAG-PRDM16 in HEK293T cells. (J) Co-IP of DNMT1 and various fragments of PRDM16. HA-tagged fragments of PRDM16 were expressed along with full-length DNMT1 in HEK293T cells. Cell lysates were immunoprecipitated with anti-DNMT1 antibodies followed by immunoblotting with HA or DNMT1 antibodies. Color-coded domain architecture of PRDM16 shows a PR/SET domain (PR), an N-terminal zinc-finger domain containing seven C2H2 zinc finger motifs (ZF1), a proline rich domain (PRR), a repression domain (RD), a second C-terminal zinc-finger domain containing three C2H2 zinc finger motifs (ZF2), and an acidic activation domain (AD). (K) Co-IP of PRDM16 and various fragments of DNMT1. HA-tagged fragments of DNMT1 were expressed along with full-length PRDM16 in HEK293T cells. Cell lysates were immunoprecipitated with anti-HA antibodies followed by immunoblotting with HA or PRDM16 antibodies. Color-coded domain architecture of DNMT1 shows the N-terminal independently folded domain (NTD), replication foci-targeting sequence (RFTS) domain, a Zn-finger like CXXC motif, two bromo adjacent homology (BAH1 and BAH2) domains, and the catalytic domain. (L) Schematic illustration of the interaction between UTX-regulated PRDM16 and DNMT1 in the maintenance of brown fat identity and suppression of myogenic remodeling in mature brown adipocytes. In brief, in mature brown adipocytes, UTX maintains the persistent demethylation of the repressive mark H3K27me3 at *Prdm16* promoter, leading to high expression of *Prdm16*; PRDM16 then recruits the DNA methyltransferase DNMT1 to *Myod1* promoter, causing *Myod1* promoter hypermethylation, and suppressing *Myod1* expression. The interaction between PRDM16 and DNMT1 coordinately serves to maintain brown adipocyte identity while repressing myogenic remodeling in mature brown adipocytes, thus promoting their active brown adipocyte thermogenic function. Suppressing this interaction by HFD feeding induces brown adipocyte-to-myocyte remodeling, which limits brown adipocyte thermogenic capacity and compromises diet-induced thermogenesis, leading to the development of obesity.

We further determined whether UTX-PRDM16 axis regulates *Myod1* promoter methylation. Indeed, knockdown of *Utx* in BAT1 brown adipocytes resulted in reduced DNA methylation at a number of CpG sites at *Myod1* promoter (**Fig 7E**). Similarly, BAT1 brown adipocytes with *Prdm16* knockdown also exhibited decreased DNA methylation at *Myod1* promoter (**Fig 7F**), while overexpression of *Prdm16* did the opposite (**Fig 7G**).

To determine how PRDM16 regulates DNA methylation at *Myod1* promoter, we knocked down *Prdm16* and measured DNMT1 binding to *Myod1* promoter via ChIP assays in BAT1 brown adipocytes with or without isoproterenol stimulation. We found that stimulation of BAT1 brown adipocytes with the β-adrenergic agonist isoproterenol significantly induced DNMT1 binding to *Myod1* promoter; whereas *Prdm16* knockdown in BAT1 brown adipocytes suppressed both basal- and isoproterenol-induced DNMT1 binding to *Myod1* promoter (**Fig 7H**), suggesting that PRDM16 is required for the full ability of DNMT1 in maintaining DNA methylation at *Myod1* promoter in brown adipocytes.

To further determine whether PRDM16 directly interacts with DNMT1, we overexpressed Flag-PRDM16 and DNMT1 in HEK293T cells and conducted co-immunoprecipitation assay. Indeed, immunoprecipitation of PRDM16 with a Flag antibody pulled down DNMT1 protein (**Fig 7I**). To further determine the domains that dictate the interaction between PRDM16 and DNMT1, we expressed various truncated fragments of PRDM16 and DNMT1 to perform co-immunoprecipitation assays in HEK293T cells. PRDM16 is a transcriptional cofactor that contains six major domains, including a PR/SET domain (PR) with monomethyltransferase activity and two zinc-finger domains (ZF1/ZF2) known to interact with PGC1α (Chi and Cohen, 2016; Kajimura et al., 2008; Nishikata et al., 2003). Other domains include a proline-rich domain (PRR), a repressor domain (RD), and a C-terminal acidic domain (AD), although the function of these domains has not been fully elucidated (Chi and Cohen, 2016; Kajimura et al., 2008; Nishikata et al., 2003). Interestingly, immunoprecipitation of DNMT1 followed by immunoblotting of HA-PRDM16 fragments with a HA antibody revealed that DNMT1 protein interacted with both PR and ZF2 domains of PRDM16 (**Fig 7J**).

Meanwhile, DNMT1 also has six functional domains including the N-terminal independently folded domain (NTD) that is known to interact with various protein that regulates DNMT1 functions, replication foci-targeting sequence (RFTS) domain, a Zn-finger like CXXC motif, two bromo adjacent homology (BAH1 and BAH2) domains, and the catalytic domain (Berkyurek et al., 2014; Tajima et al., 2016). Using a similar approach, we performed immunoprecipitation of HA-DNMT1 fragments with HA antibodies followed by immunoblotting with PRDM16 antibodies, and found that PRDM16 interacted with the NTD domain of DNMT1 protein (**Fig 7K**).

## Discussion

This study is a logical extension of our prior observation that the histone demethylase UTX promotes thermogenic program in brown adipocytes in vitro (Zha et al., 2015). Here we further determined the role of *Utx* in the regulation of brown fat thermogenesis and energy metabolism *in vivo* using genetic models and interrogated the underlying mechanism. We found that mice with *Utx* deficiency in brown adipocytes had impaired BAT thermogenesis and were susceptible to diet-induced obesity. Interestingly, we discovered that *Utx*-deficient brown adipocytes not only displayed suppressed brown adipocyte thermogenic features, they also underwent a BAT-to-myocyte-like remodeling, characterized by a profound up-regulation of myogenic markers. Interestingly, the BAT-to-myocyte remodeling was only observed in brown adipocytes from UTXKO mice, where *Utx* deletion occurred in mature brown adipocytes with the use of *Ucp1-Cre*, but not in brown adipocytes from MUTXKO mice, where *Utx* was deleted in *Myf5*-expressing myoblast precursor cells with the use of *Myf5-Cre*. Thus, our data suggest that UTX may be dispensable in the BAT-skeletal muscle lineage determination during BAT development, but it is necessary in maintaining brown adipocyte identity and suppressing myogenic remodeling in mature brown adipocytes.

Intriguingly, a parallel study involving another genetic model D1KO mice with brown adipocyte-specific *Dnmt1* deletion revealed that D1KO mice strikingly mimicked UTXKO mice in their myocyte-like brown fat remodeling, as well as in metabolic phenotypes of their susceptibility to diet-induced obesity, suggesting that both UTX and DNMT1 are involved in regulating BAT-myocyte remodeling in brown adipocytes.

Brown fat and skeletal muscle share the same developmental origin and are derived from *myf5*-expressing myoblast precursors (Seale et al., 2008; Seale et al., 2007; Wang and Seale, 2016). It is well documented that PRDM16 and MYOD1 function as reciprocal master regulators to control the cell lineage determination between brown adipogenesis and myogenesis. PRDM16 promotes BAT-specific gene expression by forming transcriptional complexes with CCAAT/enhancer binding protein β (C/EBPβ), PPARγ and PGC1α in the initiation of BAT development and subsequent maintenance of its identify (Chi and Cohen, 2016); whereas MYOD1 acts downstream of MYF5 to initiate myogenic differentiation (Rawls et al., 1998; Rudnicki et al., 1993).

We found that BAT from UTXKO mice displayed up-regulated expression of *Prdm16*, and *Utx* knockdown in BAT1 brown adipocytes resulted in enriched H3K27me3 at *Prdm16* promoter. These data suggest that *Prdm16* is a downstream target of UTX by epigenetic modulation of H3K27me3 at *Prdm16* promoter. Downstream of PRDM16, it has been shown that PRDM16 interacts with various proteins to form transcriptional complexes that mediates its effects. The ZF1/ZF2 domains of PRDM16 have been shown to interact with PPARγ (Seale et al., 2008), PGC1α (Kajimura et al., 2008), C/EBPβ (Kajimura et al., 2009), the histone H3 lysine 9 (H3K9) methyltransferase euchromatic histone methyltransferase 1 (EHMT1)(Ohno et al., 2013), and mediator complex subunit 1 (MED1) (Harms et al., 2015; Iida et al., 2015) to mediate PRDM16’s effect on maintaining brown fat identity. In addition, the PLDLS motif within the RD domain of PRDM16 has been shown to interact with C-terminal-binding protein-1 (CtBP-1) and CtBP-2 to repress WAT-specific gene expression (Kajimura et al., 2008). However, the function of the N-terminal PR/SET domain in PRDM16 is not well defined. Here we found that PRDM16 directly interacted with DNMT1 through its PR and ZF2 domain to recruit DNMT1 to *Myod1* promoter, causing *Myod1* promoter DNA hypermethylation, thus suppressing its expression, underscoring the importance of the PR/SET and ZF2 domains mediating PRDM16’s effect on *Myod1* promoter DNA hypermethylation. This was further confirmed by CRISPR/gRNA-guided approach to specifically induce demethylation at *Myod1* promoter in brown adipocytes, which resulted in increased *Myod1* expression, leading to a BAT-to-myocyte remodeling process.

Our study also demonstrates that this epigenetic event that controls BAT-myocyte remodeling may also take place in the context of diet-induced obesity and regulates diet-induced thermogenesis and systemic energy metabolism. For instance, we found that HFD decreased the binding of UTX to *Prdm16* promoter region, thus increasing H3K27me3 levels at *Prdm16* promoter. This resulted in down-regulation of *Prdm16* gene expression. Reduced PRDM16 presence would decrease the recruitment of DNMT1 to *Myod1* promoter, leading to increased *Myod1* expression and initiation of BAT-to-myocyte remodeling in BAT of HFD-fed mice. These myocyte-like brown adipocytes appeared to exhibit decreased energy expenditure as evidenced by decreased oxygen consumption rate in *Utx*- or *Dnmt1*-deficient as well as HFD-fed brown adipocytes. It has been well documented that long-term HFD feeding causes WAT remodeling, as evidenced by enlarged adipocytes, exaggerated inflammation, inappropriate fibrosis, and impaired angiogenesis (Crewe et al., 2017; Sun et al., 2011), leading to WAT dysfunction. However, much less is known about the impact of long term HFD feeding on brown adipocytes. Here we show that long-term HFD causes brown fat remodeling into a myocyte-like characteristic. The induction of these myocyte-like brown adipocytes may reduce the thermogenic capacity of brown fat, leading to reduced energy expenditure and obesity.

How HFD induces such epigenetic alterations that lead to BAT-to-myocyte remodeling in BAT is not clear. UTX catalyzes the removal of the repressive chromatin mark H3K27me3 (Ge, 2012), and H3K27 methylation has been linked to the control of cellular differentiation (Conway et al., 2015; Manna et al., 2015). In addition, cellular oxygen level also determines cell fate, and hypoxia is important in maintaining cells at an undifferentiated, precursor-like or stem cell-like state (Mohyeldin et al., 2010). Interestingly, recent data suggest that UTX, but not its paralog KDM1 lysine (K)-specific demethylase 6B/Jumonji domain-containing 3 (KDM6B/JMJD3) serves as s cellular oxygen sensor; hypoxia and loss of UTX similarly cause persistent increase in H3K27 methylation in key regulators of cellular differentiation, which in turn blocks cellular differentiation (Chakraborty et al., 2019). Hypoxia develops in obese white adipose tissue with the expansion of tissue mass, and contributes to adipose tissue dysfunction, including inflammation, fibrosis and insulin resistance (Trayhurn, 2013). Recent data also suggest that obesity induces capillary rarefaction and functional hypoxia in BAT, leading to a BAT “whitening” phenotype (Shimizu et al., 2014). Thus, it is possible that hypoxic condition induced in obese BAT may inhibit UTX activity, causing a persistent increase in H3K27 methylation at promoters of key regulators of differentiation. This in turn reverses these brown adipocytes to a less-differentiated, precursor-like phenotype, leading to a loss of brown adipocyte thermogenic feature and simultaneous up-regulation of myogenic gene expression in *Utx*-deficient brown adipocytes, as observed in the current study.

In summary, we have identified a novel BAT-myocyte remodeling process that appears to be present in mature brown adipocytes; this process is regulated by the interaction of epigenetic pathways that regulate histone and DNA methylation (**Fig 7L**). In brief, in mature brown adipocytes, UTX maintains the persistent demethylation of the repressive mark H3K27me3 at *Prdm16* promoter, leading to high expression of Prdm16. PRDM16 then recruits the DNA methyltransferase DNMT1 to *Myod1* promoter, causing *Myod1* promoter hypermethylation, and suppressing *Myod1* expression in mature brown adipocytes. The interaction between PRDM16 and DNMT1 coordinately serves to maintain brown adipocyte identity while repressing myogenic remodeling in mature brown adipocytes, thus promoting their active brown adipocyte thermogenic function. The suppression of this pathway by HFD feeding results in the induction of BAT-to-myocyte remodeling, which could limit the thermogenic capacity of brown fat and thereby compromise diet-induced thermogenesis, leading to the development of obesity (**Fig 7L**).

## Acknowledgements

This work is supported by NIH grants R01DK107544, R01DK118106 and R01DK125081, and American Diabetes Association (ADA) grant 1-18-IBS-260to BX; NIH grants R01DK115740 and R01DK118106, and ADA grant 1-18-19 IBS-348 to HS; NIH grant R01DK116496 and ADA grant 1-18-IBS-346 to LY.

## Author contribution

FL performed most of the experiments and data analysis; JJ assisted FL in these experiments; MM performed metabolic cage studies in wild type C57BL/6J mice on chow or HFD; XC, QC, RW assisted in various experiments; ZC, YP and HDS performed bioinformatics analysis of RNAseq and RRBS data; LY contributed to study design, technical inputs and review/edits on manuscript; BX and HS conceived and designed study and wrote the manuscript.

## Methods

### Mice

Mice with brown adipocyte-specific *Utx* knockout were generated by crossing *Utx*-floxed mice (Jackson Lab, stock No. 021926) (Welstead et al., 2012) with *Ucp1-Cre* mice (Jackson Laboratory, Stock No. 024670)(Kong et al., 2014), where *Ucp1* is specifically expressed in brown adipocytes and UCP1-positive beige adipocytes (UTXKO, *Ucp1^Cre^*::*Utx*^*fl*/y^ for male and *Ucp1^Cre^::Utx*^fl/fl^ for female). The littermates *Utx*^fl/Y^ (fl/Y) or *Utx*^fl/fl^ (fl/fl) were used as male or female control mice, respectively.

Mice with brown adipocyte-specific DNA methyltransferase 1 (*Dnmt1*) knockout were generated by crossing *Dnmt1*-floxed mice (Mutant Mouse Regional Resource Centers (MMRRC, No. 014114) (Jackson-Grusby et al., 2001) with *Ucp1-Cre* mice (Kong et al., 2014) (D1KO, *Ucp1^Cre^::Dnmt1*^fl/fl^), with littermates *Dnmt1*^fl/fl^ (fl/fl) as control mice.

Mice with brown adipocyte-specific *Dnmt3a* knockout were generated by crossing *Dnmt3a*-floxed mice (MMRRC No. 029885) (Kaneda et al., 2004) with *Ucp1-Cre* mice (Kong et al., 2014)(D3aKO, *Ucp1^Cre^::Dnmt3a*^fl/fl^), with littermates *Dnmt3a*^fl/fl^ (fl/fl) as control mice.

Mice with *Utx* deletion in *Myf5*-expressing cells were generated by crossing *Utx*-floxed mice with *Myf5-Cre* mice (Tallquist et al., 2000) (Jackson Laboratory, Stock No. 007893) (MUTXKO, *Myf5^Cre^*::*Utx*^*fl*/y^ for male and *Myf5^Cre^::Utx*^fl/fl^ for female), with littermates *Utx*^fl/Y^ (fl/Y) or *Utx*^fl/fl^ (fl/fl) as male or female control mice, respectively.

D1KO mice with brown adipocytes-specific GFP labeling were generated by triple-crossing *Dnmt1*-floxed mice, *Rosa-Gfp* mice (Jackson Laboratory, Stock No. 006148) (Srinivas et al., 2001) and *Ucp1-Cre* mice (D1KO-GFP, *Ucp1^Cre^*::*Dnmt1*^fl/fl^::*Rosa-Gfp*^fl/fl^), with littermates *Dnmt1*^fl/fl^::*Rosa-Gfp*^fl/fl^ as control mice (fl/fl-GFP).

C57BL/6J (B6) mice (Jackson Laboratories) were used in some experiments.

### Metabolic measurement

All animal procedures were approved by the Institutional Animal Care and Use Committee at Georgia State University. Mice were housed in a temperature- and humidity-controlled animal facility with a 12/12 h light–dark cycle and had free access to water and food. C57BL//6J wild type mice, UTXKO, D1KO, D3aKO mice and their respective littermate controls were weaned onto a regular chow diet (LabDiet 5001, LabDiet, St. Louis, MO, 13.5% calories from fat) or put on a high fat diet (HFD) (Research Diets D12492, 60% calorie from fat) diet when they were 5-6 weeks of age. In some of the experiments, a low fat diet (LFD) (Research Diets D12450B, 10% calorie from fat) was used as a control diet. Various metabolic measurements were conducted as follows. **1)** Body weight were measured weekly. **2)** Energy expenditure and locomotor activity were measured using PhenoMaster metabolic cage systems (TSE Systems, Chesterfield, MO). Food intake was measured in single-housed animals over seven consecutive days. **3)** Body composition representing fat and lean mass was analyzed using a Minispec NMR body composition analyzer (Bruker BioSpin Corporation; Billerica, MA). **4)** Fed and fasting glucose was measured by OneTouch Ultra Glucose meter (LifeScan, Milpitas, CA). Glucose and insulin sensitivity was determined by glucose tolerance and insulin tolerance tests (GTT and ITT, respectively) as we previously described (Wang et al., 2016). At the end of studies, various tissues including BAT and WAT were collected for further analysis of brown fat/beige adipocyte thermogenic program including gene expression, protein expression and immunohistochemistry.

### Cold exposure

UTXKO, D1KO mice and their respective littermate controls underwent a cold challenge (5°C) for 7 days. At the end of experiment, adipose tissues were dissected for further analysis of brown fat/beige adipocyte thermogenic gene expression, protein expression and immunohistochemistry. In some cold exposure experiments, a temperature transponder (BioMedic Data Systems, Seaford, DE) was implanted into mouse peritoneal cavity to monitor the body temperature as we previously described (Nguyen et al., 2017).

### Quantitative RT-PCR

Total RNA from tissues or cells was extracted using Tri Reagent kit (Molecular Research Center, Cincinnati, OH)(Wang et al., 2016). The mRNA of genes of interest was measured by an one-step quantitative RT-PCR with a TaqMan Universal PCR Master Mix kit (ThermoFisher Scientific, Waltham, MA) using an Applied Biosystems QuantStudio 3 real-time PCR system (ThermoFisher Scientific) as we previously described (Wang et al., 2016), and was further normalized to the housekeeping gene cyclophilin. The TaqMan primers/probes for all the genes either purchased from Applied Biosystems (ThermoFisher Scientific) or commercially synthesized were listed in **Supplemental Tables 1 and 2**.

### Immunoblotting

Protein levels of gene of interest in adipose tissue were assessed by immunoblotting as we described (Li et al., 2016; Wang et al., 2016; Zha et al., 2015). Tissues were homogenized in a modified radioimmunoprecipitation assay (RIPA) lysis buffer supplemented with 1% protease inhibitor mixture and 1% phosphatase inhibitor mixture (Sigma-Aldrich, St. Louis, MO) and tissue lysates were resolved by SDS-PAGE. Proteins on the gels were transferred to nitrocellulose membranes (Bio-Rad, Hercules, CA), which were then blocked, washed, and incubated with various primary antibodies, followed by Alexa Fluor 680-conjugated secondary antibodies (Life Science Technologies). The blots were developed with a Li-COR Imager System (Li-COR Biosciences, Lincoln, NE). The antibodies were listed in **Supplemental Table 3**.

### Immunohistochemistry (IHC)

WAT or BAT tissues were fixed in 10% neutral formalin and embedded in paraffin, which was further cut into 5 μm sections. The sections were either processed for hematoxylin and eosin (H&E) staining or immuno-staining with various antibodies as we previously described (Cao et al., 2019; Nguyen et al., 2017). The primary antibodies and the secondary antibodies were listed in **Supplemental Table 3**.

Briefly, to detect myocyte fibers in tissue sections, paraffin-embedded sections were incubated with anti-myosin heavy chain (MyHC) antibodies (MF20, Developmental Studies Hybridoma Bank (DSHB), University of Iowa, Iowa City, IA) (Seale et al., 2008) overnight at 4°C and then incubated with anti-mouse secondary antibodies labeled with Alexa fluor 488 (Invitrogen) for 1 hour at room temperature and counterstained with 4′,6-diamidino-2-phenylindole (DAPI). For in vitro study, cell cultures were fixed in 10% formalin. Fixed cells were incubated with anti-MyHC antibodies (MF20) overnight at 4°C and then incubated with anti-mouse secondary antibodies labeled with Alexa fluor 594 (Invitrogen) for 1 hour at room temperature and counterstained with DAPI. For IHC, paraffin-embedded sections of iBAT tissues were incubated with anti-UCP1 antibodies (abcam 10983) overnight at 4°C and then incubated with biotin-conjugated anti-rabbit secondary antibody (Jackson ImmunoResearch, 711-065-152) for 30 min at room temperature. The sections were washed in PBS and incubated with streptavidin-conjugated horseradish Peroxidase (VECTASTAIN® ABC Kit, PK-6100). Then the sections were washed in PBS and incubated with 3, 3-diaminobenzidine (DAB). Histology images were captured using Nikon Eclipse E800 Microscopy.

### Cell culture, SiRNA knockdown, overexpression, and oil red O staining

Immortalized brown fat preadipocytes BAT1 (obtained from Dr. Patrick Seale, University of Pennsylvania)(Rajakumari et al., 2013; Seale et al., 2007) were maintained in growth medium (DMEM/F12 containing 10% fetal bovine serum and 1% penicillin/streptomycin) at 37°C with 5% CO. The cells were differentiated into brown adipocytes as described (Rajakumari et al., 2013; Zha et al., 2015). Briefly, to differentiate brown preadipocytes, 90% confluent cells were cultured in induction medium (growth medium supplemented with 20nM insulin, 1nM T3, 125μM indomethacin, 500μM isobutylmethylxanthine (IBMX) and 0.5μM dexamethasone) for two days. After two days, cells were cultured in maintenance medium (growth medium supplemented with 20nM insulin and 1nM T3) until experiment.

For SiRNA knockdown assays, targeting siRNA and non-targeting scramble control siRNA were purchased from GE Healthcare (mouse *Dnmt1* siRNA-SMARTpool (L-056796-01), *Myod1* siRNA-SMARTpool (L-041113-00), *Prdm16* siRNA-SMARTpool (L-041318-01), and non-Targeting Scramble Control siRNA (D-001810-01)).

For plasmids overexpression and sub-cloning, full-length *Prdm16* overexpressing plasmid with an in-frame N-terminal FLAG tag was purchased from Addgene (Addgene #15504). Full-length *Dnmt1* cDNA clone was obtained from Open Biosystems and further subcloned into the pLVX lentiviral expression vector (Clontech) as we previously described (Wang et al., 2016). For sub-cloning of *Prdm16* and *Dnmt1* fragments for co-immunoprecipitation experiments, Fragments of *Prdm16* (1–223, 224–454, 455–680, 681–880, 881–1038, and 1039–1176) were PCR-amplified using the full-length Prdm16 plasmids (Addgene #15503) and sub-cloned into XbaI/EcoRI sites of Flag-HA-pcDNA3.1 vector (Addgene #52535). Fragments of *Dnmt1* (2–290, 291–603, 604–1124, 1125–1610) were PCR-amplified using the full-length pLVX-Dnmt1 plasmids and sub-cloned into XbaI/HindIII sites of Flag-HA-pcDNA3.1 vector (Addgene #52535).

SiRNAs or overexpressing plasmids were transfected into day 4 differentiated BAT1 brown adipocytes using Amaxa Nucleofector II Electroporator (Lonza) with an Amaxa cell line nucleofector kit L according to the manufacturer’s instructions (Lonza) as we previously described (Li et al., 2016; Zha et al., 2015). Briefly, at days 4 of differentiation, cells (2 × 10^6^ cells/sample) were trypsinized and centrifuged at 90 × g for 5 min at room temperature. Cells were then resuspended in nucleofector solution (100 μl/sample) with 20 pmol of different siRNAs or plasmids and seeded into 24-well plates or 100 mm dishes. Cells were harvested 2 days after for further analysis.

For oil red O staining assays, differentiated BAT1 cells were fixed with 10% formalin and washed twice with ddH2O. Cells were then incubated with 0.05% oil red O (Sigma: 00625-25G) working solution as we described previously (Chen et al., 2016). Samples were visualized using Nikon Eclipse E800 Microscopy.

### Measurement of oxygen consumption in brown adipocytes

Cellular oxygen consumption in brown adipocytes was measured using a XF 96 Extracellular Flux Analyzer (Agilent, Santa Clara, CA) as we previously described (Shin et al., 2017). The measurement started with basal respiration recording followed by addition of a sequential reagents including oligomycin for inhibition of the coupled respiration and FCCP for maximal respiration.

### RNA-sequencing analysis

Total RNA was isolated from iBAT as described above and was submitted to Beijing Genomics Institute (BGI, Shenzhen, China) for RNA-sequencing (RNAseq) analysis. Equal amount of RNAs from 6 animals/group were pooled and used for RNAseq analysis. Clean reads were aligned to the mouse reference genome (UCSC mm9). Differentially expressed genes between groups were defined as Log2 fold change ≥0.5 or ≤-0.5. The RNAseq data was also used to predict adipose tissue browning capacity with an online bioinformatic software https://github.com/PerocchiLab/ProFAT (Cheng et al., 2018).

### Whole genome DNA methylation analysis with reduced representation bisulfite sequencing (RRBS) to identify Myod1 promoter methylation

Genomic DNA from BAT was isolated by phenol chloroform extraction and a commercial service for DNA methylation sequencing was provided by Beijing Genomics Institute (BGI) (Shenzhen, China). Equal amount of genomic DNAs from 6 animals/group were pooled and used for RRBS analysis. According to the brochure provided by the company, the genomic DNA was digested with the methylation-insensitive restriction enzyme MspI and ligated to methylated sequencing adaptors. The ligated MSPI fragments were size-selected, treated with sodium bisulfite, amplified by PCR and constructed for library, which was sequenced. Clean reads were aligned to genome annotation datasets (University of California Santa Cruz (UCSC) Genome Browser on Mouse (NCBI37/mm9) Assembly). The software tool Bioconductor, a software repository of R packages, was used to produce such annotations. The tool kit methylKit v0.9.6. was used to generate methylation report for each sample, and Log2 fold change threshold of 0.5 was used to identify Differentially Methylated Regions (DMRs) between two genotypes.

### Chromatin immunoprecipitation (ChIP) assays

ChIP assays were conducted using a ChIP assay kit (Upstate) as we previously described (Li et al., 2016; Zha et al., 2015). Tissues were fixed and used for nuclei isolation. The nuclei were resuspended in lysis buffer and sonicated to shear DNA, followed by immunoprecipitation, elution, and analyzed by real-time PCR using SYBR green. Primer sequences used in this study were: *Myod1*, forward: 5’-ACTCCTATTGGCTTGAGGCG-3’, reverse: 5’-CAAGCCGTGAGAGTCGTCTT-3’; *Prdm16*, forward: 5’-ACGAAGAGGATGATGAACACATT-3’, reverse: 5’-TCATCTCCCTAGCATTGTCAGTT-3’.

### Bisulfite conversion and pyrosequencing

The detection of DNA methylation levels at *Myod1* promoter and 5’-region was conducted as we previously described (Wang et al., 2016). Genomic DNA was isolated by phenol/chloroform extraction, followed by bisulfite conversion with an EpiTech Bisulfite Kit (Qiagen). The primers for PCR-amplification of *Myod1* proximal promoter/5’-region and for sequencing are shown in **Supplemental Table 4**. Bisulfite-converted DNA (2μg) was amplified by PCR, which was sent to EpiGenDx (Hopkinton, MA) for sequencing analysis.

### Targeted demethylation at the Myod1 promoter

A endonuclease dead version of Cas9 (dCas9) has been engineered to be fused with the catalytic domain of the enzyme involved in DNA demethylation, TET1 (dCas9-TET1CD) (Liu et al., 2016; Vojta et al., 2016). The mammalian lentiviral vectors FUW carrying dCas9-TET1CD were purchased from Addgene (Addgene No. 84475). Guide sequences targeting the CpG sites at the *Myod1* promoter was designed with GT-Scan website (http://gt-scan.braembl.org.au/gt-scan) and targeting or non-targeting oligos were annealed and inserted into the AarI sites of the pgRNA lentiviral vector that co-express mCherry (Addgene No. 44248). The gRNA sequence for *Myod1*-targeting oligo was: forward, 5’-caccGTACTGTTGGGGTTCCGGAGTGG-3’, reverse, 5’-aaacCCACTCCGGAACCCCAACAGTAC-3’. The sequence for non-targeting gRNA was: forward, 5’-ttggCCCCCGGGGGAAAAATTTTT; reverse, 5’-aaacAAAAATTTTTCCCCCGGGGG-3’ (Liu et al., 2016). Lentiviruses expressing dCas9-TET1CD or gRNA-mCherry (1×10^9^IFU/ml) was produced by Vigene Biosciences, Inc., and was bilaterally injected into iBAT according to previously published methods (Balkow et al., 2016; Liu et al., 2016; Nguyen et al., 2017). Briefly, a small skin incision was made above iBAT, and a series of microinjections with designated lentivirus (1×10^9^IFU/ml) were given across five evenly distributed loci (2μl/locus) for each iBAT pad (10 total injections per animal) and the needle left in place for 1 minute following each injection to prevent efflux. After the final injection, the skin was closed with sterile wound clip staples and animals were returned to their cages. Animals were kept isolated in their home cage for 7 days for recovery.

### Statistical analysis

Data were expressed as mean ±SEM. Statistical tests were performed using SPSS software (version 16.0, SPSS Inc, Chicago, IL, USA). Experimental data were analyzed by unpaired Student’s t-test or one-way Analysis of Variance (ANOVA) followed by Bonferroni post-hoc analysis where appropriate. Statistical significance was accepted at *p*<0.05.

