## Supplemental Tables and Figures for "Epigenetic Interaction between UTX and DNMT1 Regulates Diet-Induced Myogenic Remodeling in Brown Fat"

**Running title**: UTX and DNMT1 regulate myogenic remodeling in BAT.

* Correspondence should be addressed to:

Hang Shi, Department of Biology and Center for Obesity Reversal, Georgia State University, Atlanta, GA 30303, USA. Contact: 404-413-5799,.

Bingzhong Xue, Department of Biology and Center for Obesity Reversal, Georgia State University, Atlanta, GA 30303, USA. Contact: 404-413-5747,

**Supplemental Table 1. Primer/probe sets for gene expression**

| Gene symbol | Company | Catalog # |
|---|---|---|
| ***Acox1*** | ABI | Mm01246834_m1 |
| ***Acta1*** | ABI | Mm00808218_g1 |
| ***Atp2a1*** | ABI | Mm01275320_m1 |
| ***Cd137*** | ABI | Mm01268456-m1 |
| ***Cidea*** | ABI | Mm00432554_m1 |
| ***Ckm*** | ABI | Mm01321487_m1 |
| ***Cox*** | ABI | Mm04225243_g1 |
| ***Cpt1b*** | ABI | Mm00487191_g1 |
| ***Dio2*** | ABI | Mm0051664_m1 |
| ***Dnmt1*** | ABI | Mm00599783-g1 |
| ***Ear2*** | ABI | Mm04207376_gH |
| ***Ebf2*** | ABI | Mm00438625_m1 |
| ***Ebf3*** | ABI | Mm00438642_m1 |
| ***Elovl3*** | ABI | Mm01194165_g1 |
| ***Eva1*** | ABI | Mm00468397_m1 |
| ***Fgf21*** | ABI | Mm00840165_g1 |
| ***Klf2*** | ABI | Mm00500486_g1 |
| ***Klhl13*** | ABI | Mm00470674_m1 |
| ***Mfsd2a*** | ABI | Mm01192208_m1 |
| ***Myh1*** | ABI | Mm01332489_m1 |
| ***Myh4*** | ABI | Mm01332541_m1 |
| ***Myod1*** | ABI | Mm00440387_m1 |
| ***Myog*** | ABI | Mm00446194_m1 |
| ***Nr4a1*** | ABI | Mm00440945-m1 |
| ***Nr4a2*** | ABI | Mm00443060_m1 |
| ***Nr4a3*** | ABI | Mm00450071_g1 |
| ***Otop1*** | ABI | Mm00554705_m1 |
| ***Pparγ*** | ABI | Mm00440945_m1 |
| ***Prdm16*** | ABI | Mm00712556_m1 |
| ***Sik1*** | ABI | Mm00440317_m1 |
| ***Ttn*** | ABI | Mm00621005_m1 |
| ***Utx*** | ABI | Mm01283053_m1 |

**Table 2. Primer/probe sequences for gene expression**

| Gene | Primer (Forward: 5'-3') | Primer (Reverse: 5'-3') | Probe (5'-3') |
|---|---|---|---|
| **Cyclophilin (*Ppib*)** | GGTGGAGAGCACCAAGACAGA | GCCGGAGTCGACAATGATG | ATCCTTCAGTGGCTTGTCCCGGCT |
| ***Cebpβ*** | CCAAGAAGACGGTGGACAAG | GTGCTGCGTCTCCAGGTT | CCGCATCTTGTACTCGTCGCTCAG |
| ***Pgc1α*** | CATTTGATGCACTGACAGATGGA | CCGTCAGGCATGGAGGAA | CCGTGACCACTGACAACGAGGCC |
| ***Pgc1β*** | AGGAAGCGGCGGGAAA | CTACAATCTCACCGAACACCTCAA | AGAGATTTCGAATGTATACCACACGGCCTTCA |
| ***Ucp1*** | CACCTTCCCGCTGGACACT | CCCTAGGACACCTTTATACCTAATGG | AGCCTGGCCTTCACCTTGGATCTGA |

**Supplemental Table 3. Antibodies used in Immunoblotting and ChIP-qPCR**

| Antibody | Company | Catalog # | Application |
|---|---|---|---|
| **DNMT1** | abcam | Ab87654 | WB |
| **DNMT1** | Santa Cruz | Sc20701 | ChIP |
| **GFP** | Aves labs | GFP-1010 | IF |
| **UCP1** | abcam | ab23841 | WB |
| **UCP1** | abcam | Ab10983 | IHC |
| **KDM6A (UTX)** | abcam | ab36938 | ChIP |
| **KDM6A (UTX)** | Bethyl Laboratories | A302-374A | WB |
| **HA** | Cell signaling technology | C29F4 | IP, WB |
| **H3K27me3** | Cell signaling technology | 9733P | ChIP |
| **mCherry** | abcam | ab205402 | IF |
| **MyHC** | DSHB | MF20 | IF |
| **Perilipin** | Everest biotech | EB07728 | IF |
| **PRDM16** | Sigma | SAB2900806 | WB, IP |
| **FLAG** | Sigma | F3165 | WB,IP |
| **α-Tubulin** | Santa Cruz | sc-53646 | WB |
| **Biotin-SP (long spacer) AffiniPure Donkey Anti-Rabbit IgG (H+L)** | Jackson ImmunoResearch | 711-065-152 | IHC |
| **Cy™3 AffiniPure Donkey Anti-Rabbit IgG (H+L)** | Jackson ImmunoResearch | 711-165-152 | IF |
| **Alexa Fluor® 488 AffiniPure Donkey Anti-Chicken IgY (IgG) (H+L)** | Jackson ImmunoResearch | 703-545-155 | IF |
| **Goat anti-Mouse IgG (H+L) Highly Cross-Adsorbed Secondary Antibody, Alexa Fluor 488** | Invitrogen | A11029 | IF |
| **Goat anti-Mouse IgG (H+L) Highly Cross-Adsorbed Secondary Antibody, Alexa Fluor 594** | Invitrogen | A11032 | IF |
| **Goat anti-Mouse IgG (H+L) Highly Cross-Adsorbed Secondary Antibody, Alexa Fluor 680** | Invitrogen | A21058 | WB |
| **Goat anti-Rabbit IgG (H+L) Highly Cross-Adsorbed Secondary Antibody, Alexa Fluor 680** | Invitrogen | A21109 | WB |

**Supplemental Table 4. Primer sequences for pyrosequencing**

| **Primers** | **Sequence (5'-3')** |
|---|---|
| *Myod1*-pyroseq-F1 | TGGTTATTTTGGGGATTTTAAGTT |
| *Myod1*-pyroseq-*R1 | TCTACCCCTCCTCCCTAT |
| *Myod1*-pyroseq-S1 | ATTTTGGGGATTTTAAGTTT |
| *Myod1*-pyroseq-F2 | GATAGGGAGGAGGGGTAGAGGATA |
| *Myod1*-pyroseq-*R2 | TCCCAATTCCTAAATCCAACCTCAAC |
| *Myod1*-pyroseq-S2 | AGGGGTAGAGGATAG |
| *Myod1*-pyroseq-F3 | GTTTGGGTTGAGGTTGGATTTA |
| *Myod1*-pyroseq-*R3 | CTCCCATTTCAAAAAAACTCCCATATACAC |
| *Myod1*-pyroseq-S3 | GGAATTGGGATATGGAG |
| *Myod1*-pyroseq-F4 | TGGTGTATATGGGAGTTTTTTTGAA |
| *Myod1*-pyroseq-*R4 | AACCTCATTCACTTTACTCAA |
| *Myod1*-pyroseq-S4 | ATGGGAGTTTTTTTGAAAT |
| *Myod1*-pyroseq-F5 | AGTTGGGTTGGATTGTTATGT |
| *Myod1*-pyroseq-*R5 | ACCACTACCCCTAATCCC |
| *Myod1*-pyroseq-S5 | ATGTAGGGTTGGAGA |
| *Myod1*-pyroseq-F6 | GGGTTAGGGATTAGGGGTAG |
| *Myod1*-pyroseq-*R6 | CCTTACTCCAAAAATCCTCAAACTC |
| *Myod1*-pyroseq-S6 | GGGATTAGGGGTAGT |

F: Forward primer; R: reverse primer; S: sequencing primer.
*Indicates primers with biotin-tag.

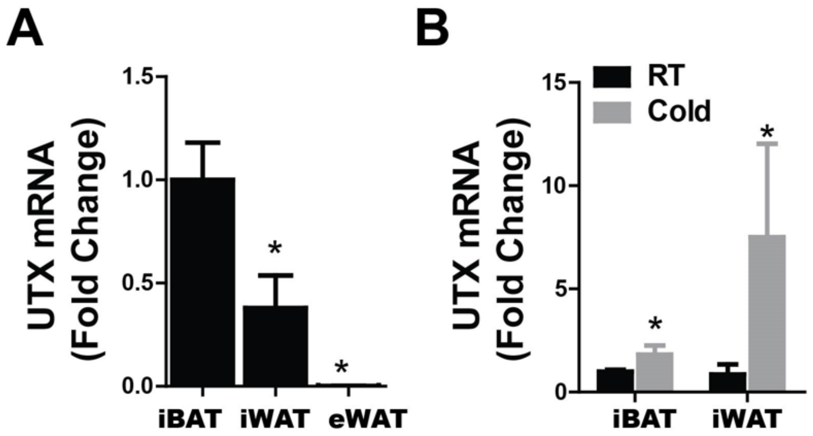


**Supplemental figure 1 (Related to Figure 1).** (A) *Utx* mRNA levels in interscapular BAT (iBAT), inguinal WAT (iWAT) and epididymal WAT (eWAT) depots of 2-month-old male mice (n=7/group). (B) *Utx* mRNA levels in iBAT and iWAT depots of 2-month-old male mice housed at room temperature (RT) or challenged with a 7-day 5°C cold exposure (n=5-6/group).

All data are expressed as mean ± SEM. *p<0.05 vs. iBAT in (A) by ANOVA with post-hoc test and RT in (B) by two-tailed unpaired Student's t-test.

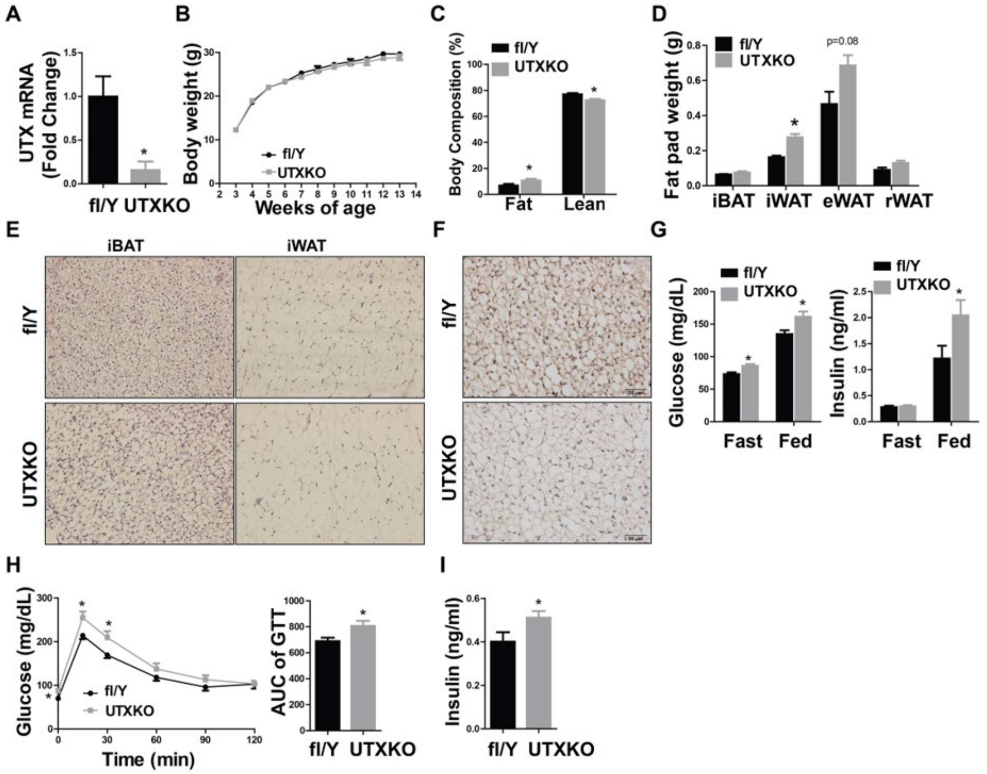

**Supplemental figure 2 (Related to Figure 1).** UTX deficiency in brown fat promotes adiposity in mice on a chow diet. Male UTXKO and their fl/Y littermates were weaned onto regular chow diet.

(A) *Utx* mRNA levels in iBAT of male UTXKO and fl/Y mice (n=7-10/group).

(B) Body weight growth curve of male UTXKO mice and fl/Y mice on regular chow diet (n=7-10/group).

(C)-(D) Body composition (C), and Fat pad weight (iBAT, iWAT, eWAT and retroperitoneal WAT (rWAT)) (D) in male UTXKO and fl/Y mice on regular chow diet (n=6-8/group).

(E)-(F) H&E staining of iBAT and iWAT (E), and UCP1 immunohistochemistry (IHC) staining in iBAT (F) in male UTXKO and fl/Y mice on regular chow diet (n=3/group).

(G)-(I) Fed and fasted circulating glucose and insulin levels (G), Glucose tolerance test (GTT) (H), and Insulin levels at 15 minutes during GTT test (I) in male UTXKO and fl/Y mice on regular chow diet (n=7-10/group).

All data are expressed as mean ± SEM. *p<0.05 vs. fl/Y by two-tailed unpaired Student’s t-test.

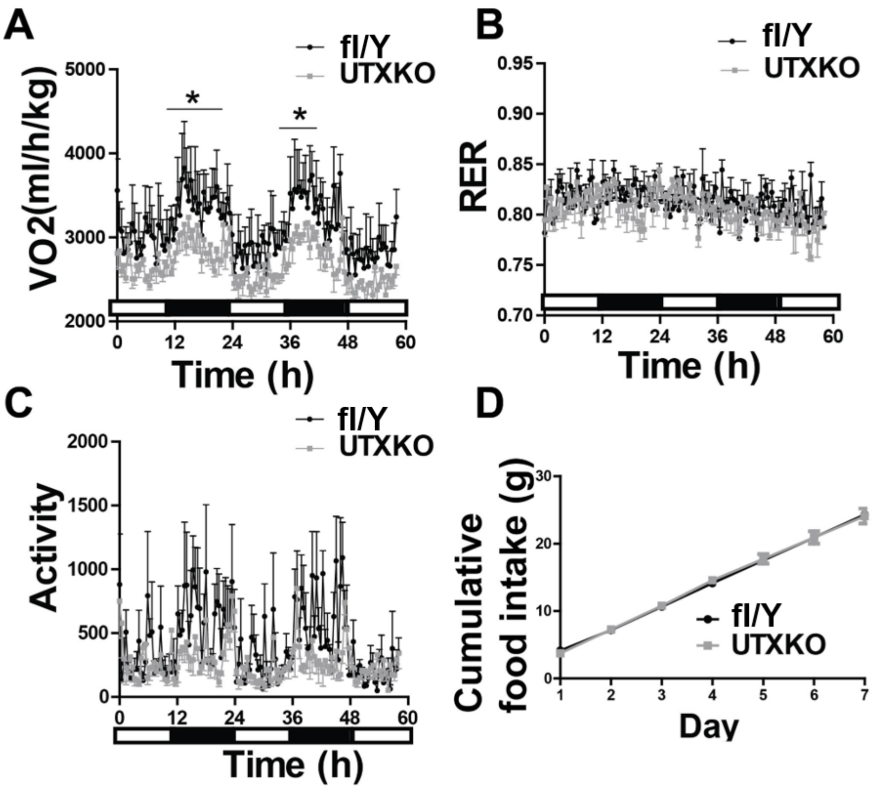


**Supplemental figure 3 (Related to Figure 1).** Metabolic characterization of male UTXKO and fl/Y control mice on HFD. Male UTXKO and their littermate control fl/Y mice were put on HFD when they were 5 weeks of age.

(A)-(D) Oxygen consumption ($VO_2$) (A), Respiratory exchange rate (RER) (B), Locomotor activity (C), and Cumulative food intake (D) in male UTXKO and fl/Y mice on HFD (n=4/group).

All data are expressed as mean ± SEM. *p<0.05 vs. fl/Y by two-tailed unpaired Student’s t-test.

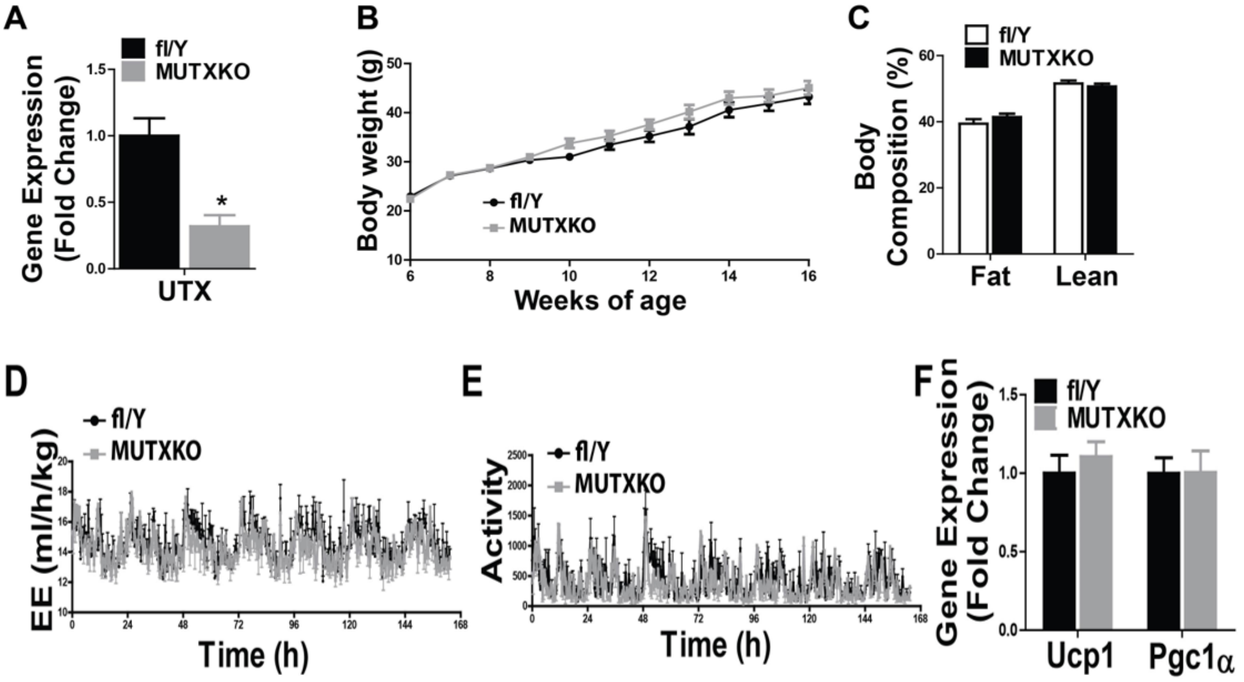


**Supplemental figure 4 (Related to Figure 1).** Mice with *Utx* deficiency in *Myf5*-expressing precursor cells (MUTXKO) have normal energy homeostasis and brown fat thermogenic gene expression when fed HFD diet. Male MUTXKO and fl/Y littermate control mice were put on HFD when they were 6 weeks of age.

(A) *Utx* mRNA levels in iBAT of male MUTXKO and fl/Y mice (n=8/group).

(B)-(C) Body weight growth curve (B), and Body composition (C) of male MUTXKO mice and fl/Y mice on HFD diet (n=9-13/group).

(D)-(E) Energy expenditure (EE)(D), and Locomotor activity (E) in male MUTXKO and fl/Y mice on HFD (n=4/group).

(F) *Ucp1* and *Pgc1α* expression in iBAT of male MUTXKO and fl/Y mice on HFD (n=8/group).

All data are expressed as mean ± SEM. *p<0.05 vs. fl/Y by two-tailed unpaired Student's t-test.

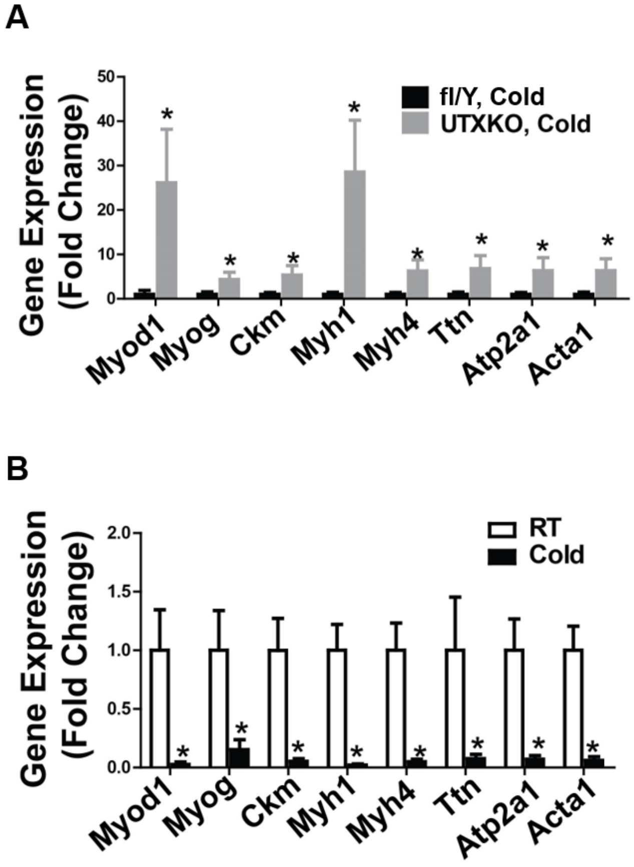


**Supplemental Figure 5 (Related to Figure 2).** Myogenic marker gene expression is up-regulated in iBAT from UTXKO mice after a 7-day 5°C cold challenge but down-regulated in iBAT from wild type C57BL/6J mice after a 7-day 5°C cold challenge.

(A) Quantitative RT-PCR analysis of myogenic marker gene expression in iBAT of chow-fed 2-month-old male UTXKO and fl/Y mice after a 7-day cold challenge (n=8/group).

(B) Quantitative RT-PCR analysis of myogenic marker gene expression in iBAT of chow-fed 2-month-old male C57BL/6J mice after a 7-day cold challenge (n=8/group).

All data are expressed as mean ± SEM. *$p<0.05$ by two-tailed unpaired Student's t-test.

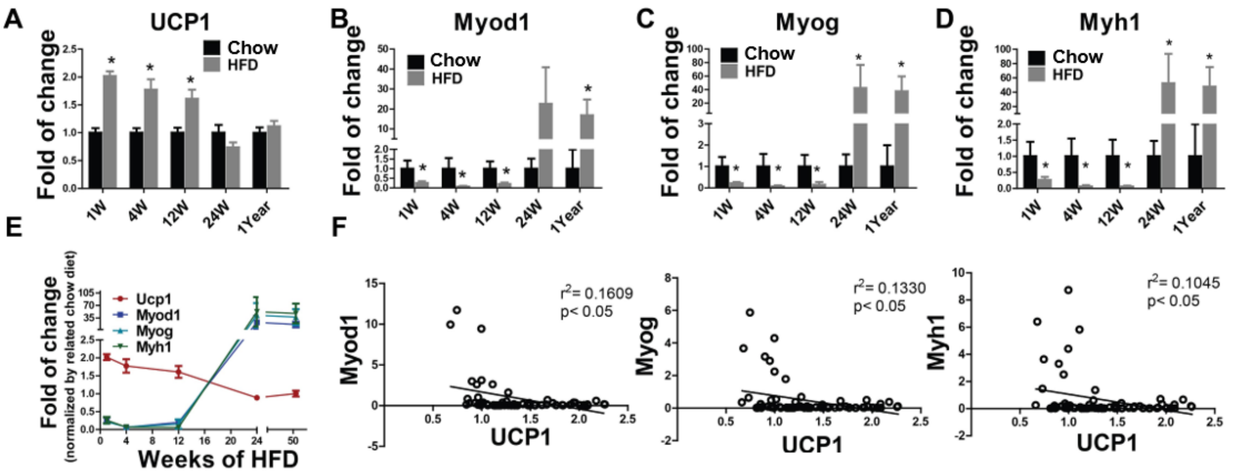


**Supplemental figure 6 (Related to Figure 2).** The expression of *Ucp1* and myogenic marker genes are negatively correlated in iBAT. Male C57BL/6J mice were put on either chow or HFD diet when they were 6 weeks old.

(A)-(D) Quantitative RT-PCR analysis of *Ucp1* (A), *Myod1* (B), *Myog* (C), and *Myh1* (D) expression in iBAT of mice fed chow or HFD for 1 week, 4 weeks, 12 weeks, 24 weeks and 1 year (n=8/group).

(E) Quantitative RT-PCR analysis of *Ucp1* and myogenic marker gene expression patterns in iBAT of HFD-fed mice for 1 week, 4 weeks, 12 weeks, 24 weeks and 1 year (n=8/group).

(F) Negative correlations between *Ucp1* and myogenic marker gene expression in iBAT of mice fed chow or HFD for 1 week, 4 weeks, 12 weeks, 24 weeks and 1 year (n=80).

All data are expressed as mean ± SEM. *p<0.05 vs. Chow by two-tailed unpaired Student's t-test in (A)-(D), two-tailed Pearson's Correlation test in (F).

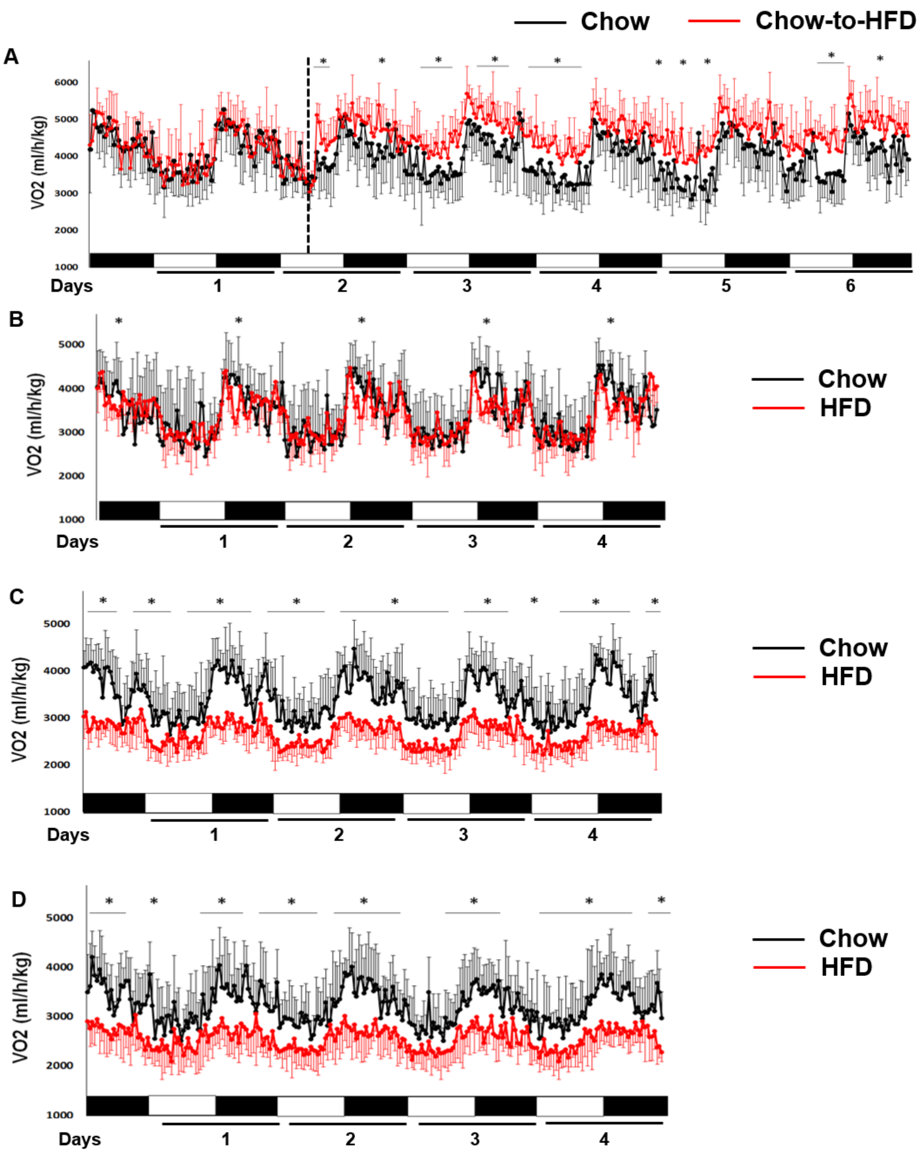


**Supplemental figure 7 (Related to Figure 2).** Oxygen consumption in wild type C57BL/6J mice fed a regular chow or HFD. Male C57BL/6J mice were put on either chow or HFD diet when they were 6 weeks old.

(A) Oxygen consumption in 6-week-old wild type C57BL/6J mice on chow diet or in 6-week-old C57BL/6J mice switching from chow to HFD. The dotted line indicates the time point when diet was switched from chow to HFD (n=8/group).

(B)-(D) Oxygen consumption in wild type C57BL/6J mice fed a regular chow or HFD for 4 weeks (B), 12 weeks (C) and 24 weeks (D) (n=8/group).

All data are expressed as mean ± SEM; (n=8/group). *$p<0.05$ vs. Chow by tow-tailed unpaired Student's t-test.

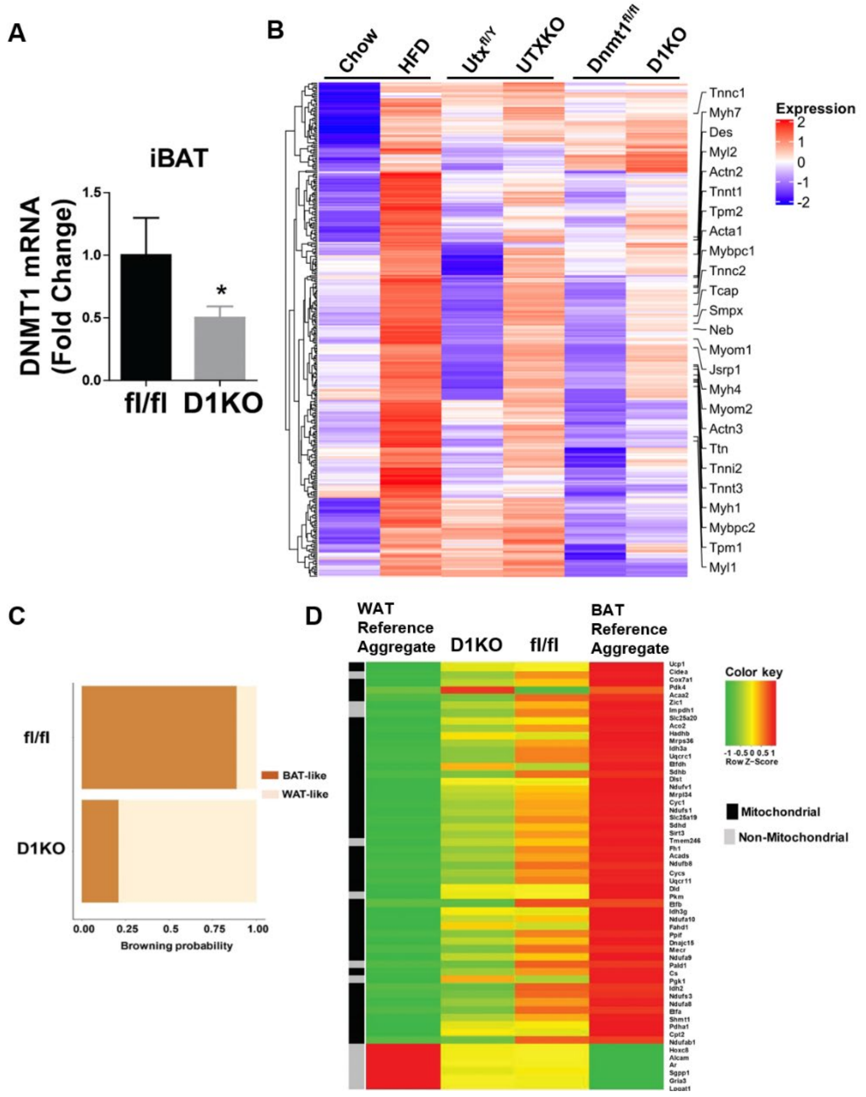


**Supplemental figure 8 (Related to Figure 3).** *Dnmt1*-deficient brown fat exhibits reduced BAT-specific gene expression and has a gene expression profile characteristic of white fat.

(A) *Dnmt1* expression in iBAT of D1KO and fl/fl mice (n=6/group).

(B) Hierarchical cluster analysis of genes similarly up-regulated in iBAT of HFD-fed, UTXKO and D1KO mice.

(C) Bioinformatic modeling of BAT-like or WAT-like gene expression profiles using RNA-seq data from iBAT of D1KO and fl/fl mice on chow diet using an online software (https://github.com/PerocchiLab/ProFAT). .

(D) BAT-specific gene expression in iBAT of D1KO and fl/fl mice on chow diet using an online software (https://github.com/PerocchiLab/ProFAT). The WAT reference aggregate and BAT reference aggregate were derived from the online software.

All data are expressed as mean ± SEM. *p<0.05 vs. fl/fl by two-tailed unpaired Student's t-test.

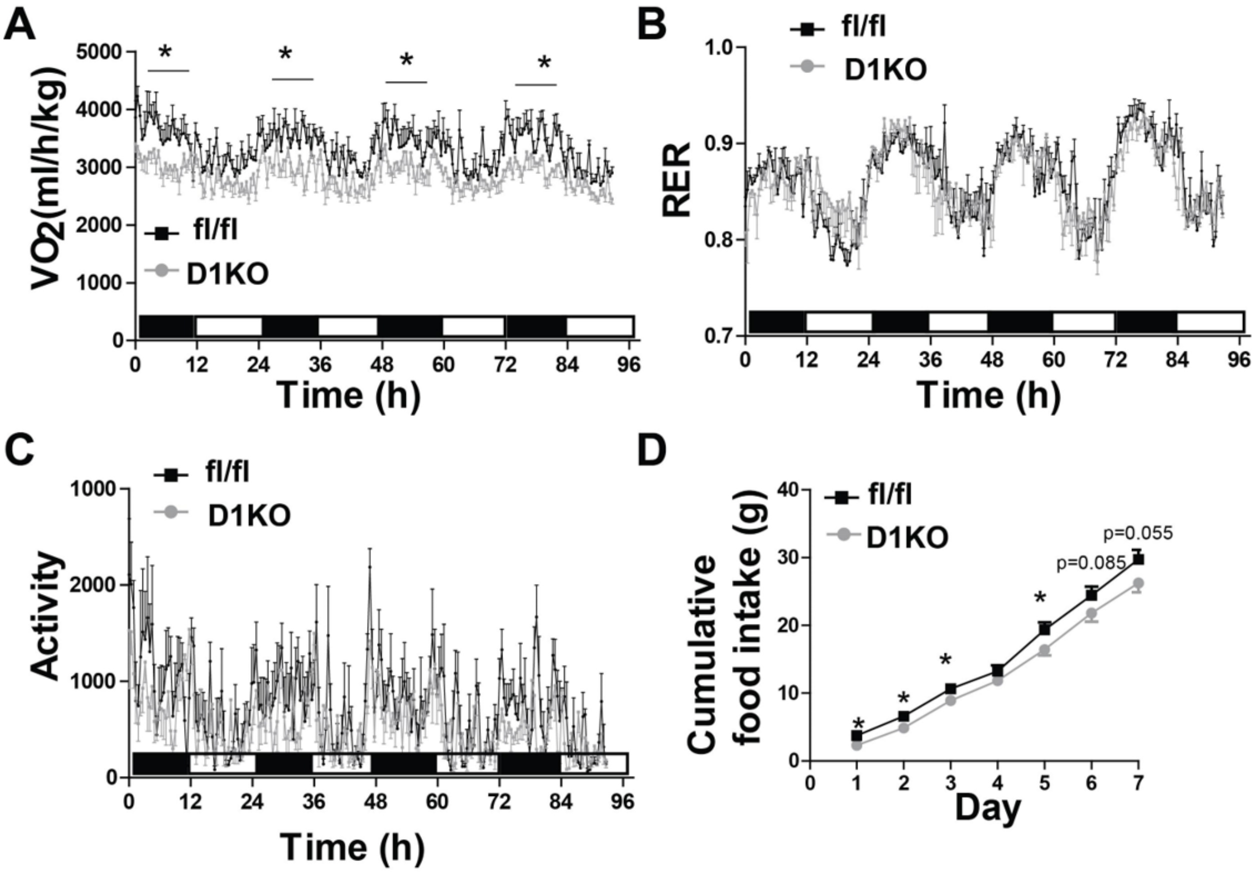


**Supplemental figure 9 (Related to Figure 4).** Metabolic characterization of female D1KO and fl/fl mice on chow diet.

(A)-(D) Oxygen consumption (A), RER (B), Locomotor activity (C), and Cumulative food intake (D) in female D1KO mice on chow diet (n=4/group).

All data are expressed as mean ± SEM. *p<0.05 by two-tailed unpaired Student's t-test.

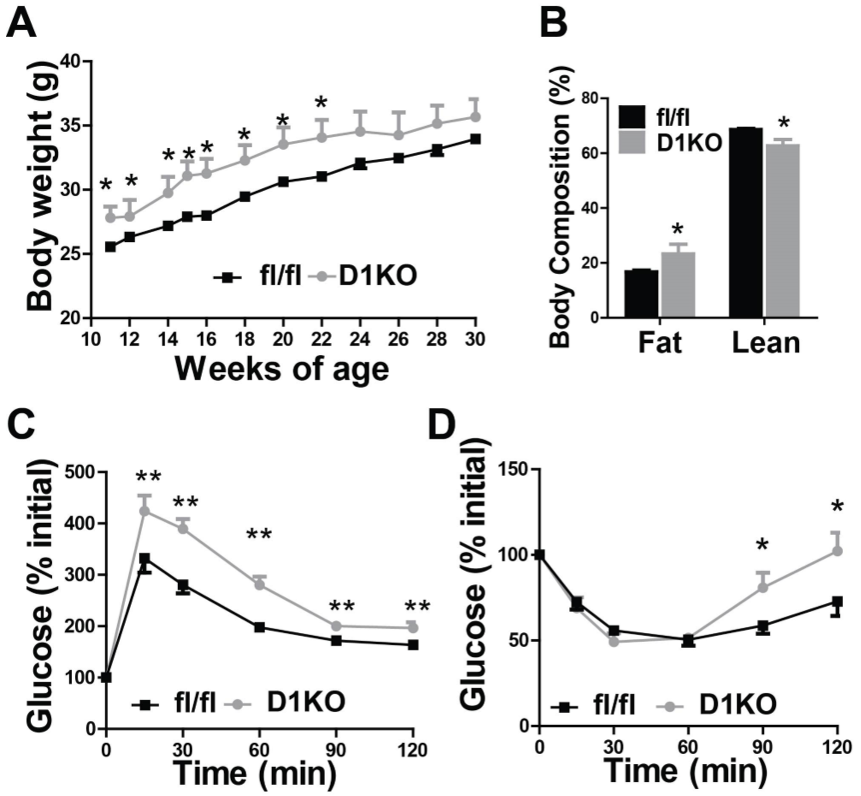


**Supplemental figure 10 (Related to Figure 4).** Metabolic characterization of male D1KO and their control fl/fl mice on chow diet.

(A)-(D) Body weight growth curve (A), Body composition (B), GTT (C), and ITT (D) in male D1KO and their control fl/fl mice on chow diet (n=8/group).

All data are expressed as mean ± SEM. *p<0.05 by two-tailed unpaired Student's t-test.

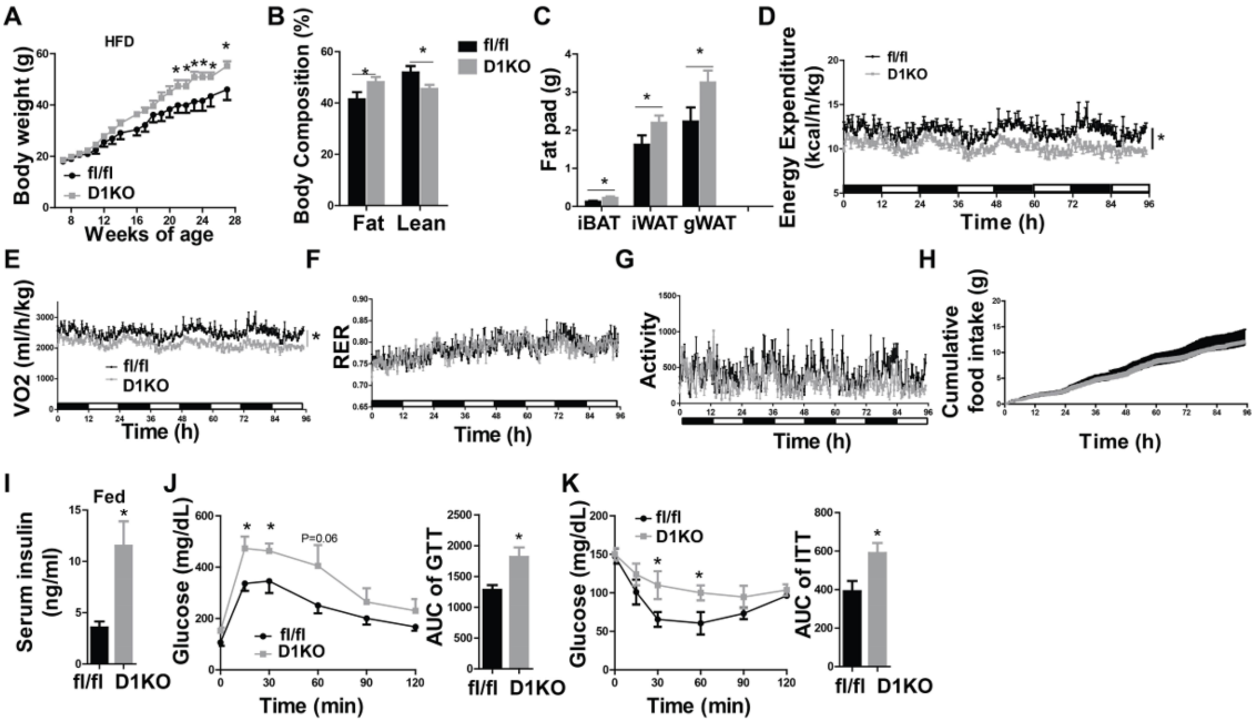


**Supplemental figure 11 (Related to Figure 4).** Female D1KO mice are susceptible to DIO. Female D1KO and fl/fl were put on HFD diet when they were 6 weeks old.

(A)-(C) Body weight growth (A), Body composition (B), and Fat pad weight (C) of female D1KO mice and their littermate fl/fl controls fed HFD (n=8/group).

(D)-(H) Energy expenditure (D), Oxygen consumption (E), RER (F), Locomotor activity (G), and Cumulative food intake (H) of female D1KO mice and their littermate fl/fl controls on HFD (n=4/group).

(I)-(K) Fed insulin levels (I), GTT (J), and ITT (K) of female D1KO mice and their littermate fl/fl controls on HFD (n=8/group).

All data are expressed as mean ± SEM. *p<0.05 by two-tailed unpaired Student's t-test.

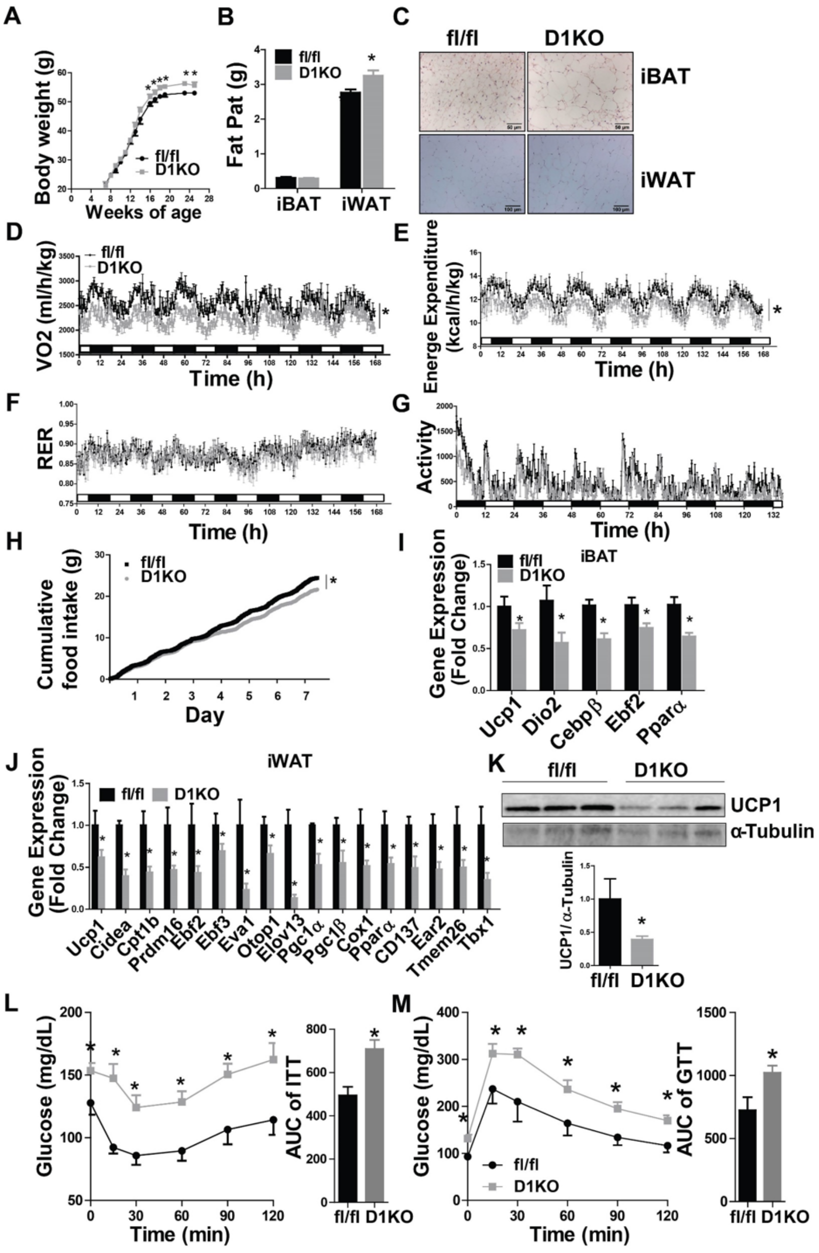


**Supplemental figure 12 (Related to Figure 4).** Metabolic characterization of male D1KO and their control fl/fl mice on HFD. Male D1KO and fl/fl mice were put on HFD diet when they were 6 weeks of age.

(A)-(B) Body weight growth curve (A) and Fat pad weight (B) in male D1KO and fl/fl mice on HFD (n=8/group).

(C) H&E staining of iBAT and iWAT in male D1KO and fl/fl mice on HFD (n=3/group).

(D)-(H) Oxygen consumption (D), Energy expenditure (E), RER (F), Locomotor activity (G), and Cumulative food intake (H) in male D1KO and fl/fl mice on HFD (n=4/group).

(I)-(J) Quantitative PCR analysis of thermogenic gene expression in iBAT (I) and iWAT (J) of male D1KO and fl/fl mice on HFD (n=8/group).

(K) Immunoblotting of UCP1 protein in iBAT of male D1KO and fl/fl mice on HFD (n=3/group).

(L)-(M) GTT (L) and ITT (M) in male D1KO and fl/fl mice on HFD (n=8/group).

All data are expressed as mean ± SEM. *$p<0.05$ by two-tailed unpaired Student's t-test.

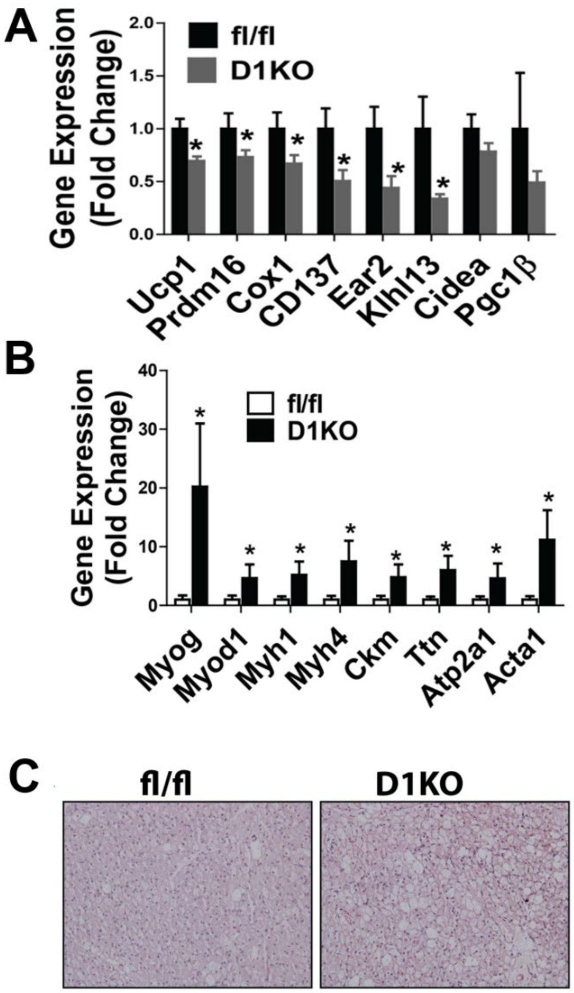


**Supplemental figure 13 (Related to Figure 4).** Characterization of D1KO and fl/fl during cold exposure. Two months old male D1KO and fl/fl mice were subjected to a chronic 7-day cold exposure challenge at 5°C.

(A)-(B) Quantitative RT-PCR analysis of thermogenic gene expression (A) and myogenic marker gene expression (B) in iBAT of male D1KO and fl/fl mice after chronic 7-day cold exposure (n=6/group).

(C) H&E staining of iBAT from male D1KO and fl/fl after a chronic 7-day cold exposure (n=3/group).

All data are expressed as mean ± SEM. *p<0.05 by two-tailed unpaired student's t-test.

Suppl. Fig. 14

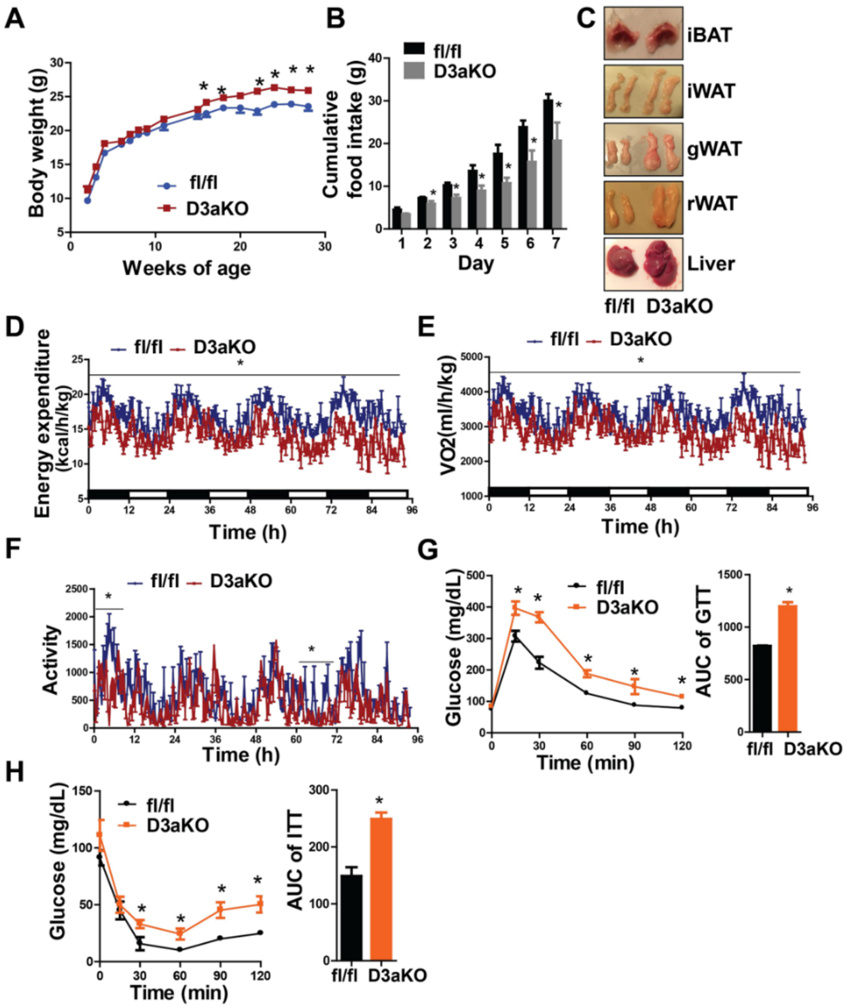


**Supplemental figure 14 (Related to Figure 4).** Metabolic characterization of D3aKO and their control fl/fl mice on chow diet.

(A)-(B) Body weight growth curve (A) and Food intake (B) in D3aKO and their control fl/fl mice on chow diet (n=4/group).

(C) Representative fat depot and liver images in D3aKO and their control fl/fl mice on chow diet (n=4/group).

(D)-(H) Energy expenditure (D), Oxygen consumption (E), Locomotor activity (F), GTT (G), and ITT (H) in D3aKO and their control fl/fl mice on chow diet (n=4/group).

All data are expressed as mean ± SEM. *p<0.05 by two-tailed unpaired Student's t-test.

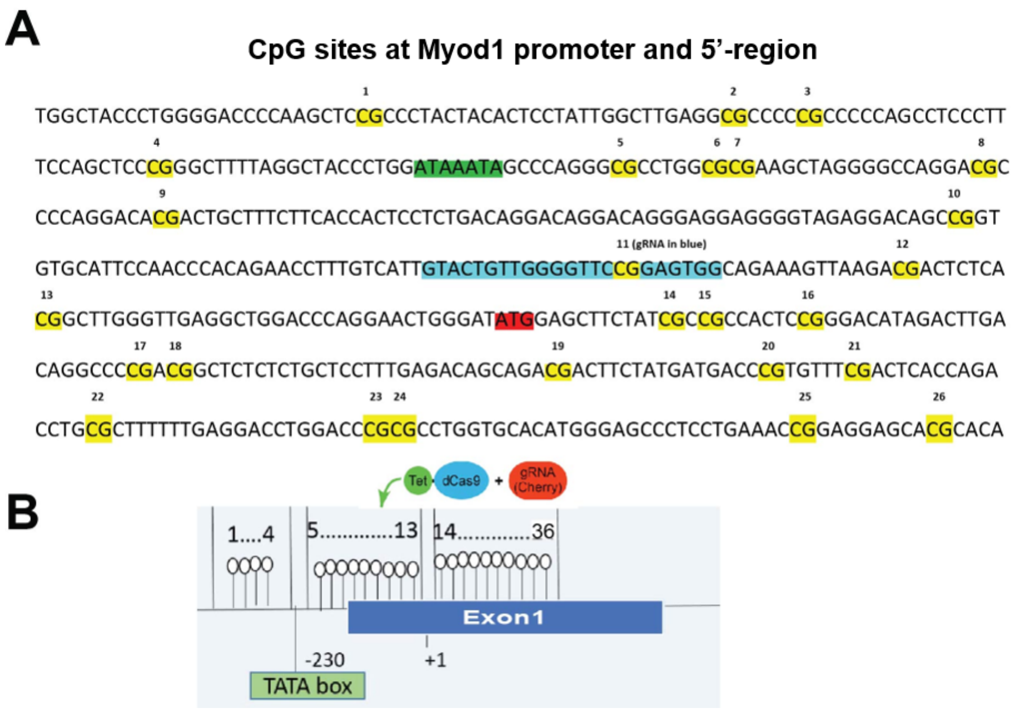


**Supplemental figure 15 (Related to Figure 5 and Figure 6).** (A) CpG sites at *Myod1* promoter and 5'-region. Yellow highlights indicate CpG sites; green highlight indicates the TATAA-box; red highlight indicates translational start site (ATG); blue-highlighted region indicates the location of guide RNA (gRNA) that was used in the directed methylation targeting approach described in (B). (B) Schematic diagram of the strategy to specifically target demethylation at *Myod1* promoter with dCas9-TET1CD and gRNA system.

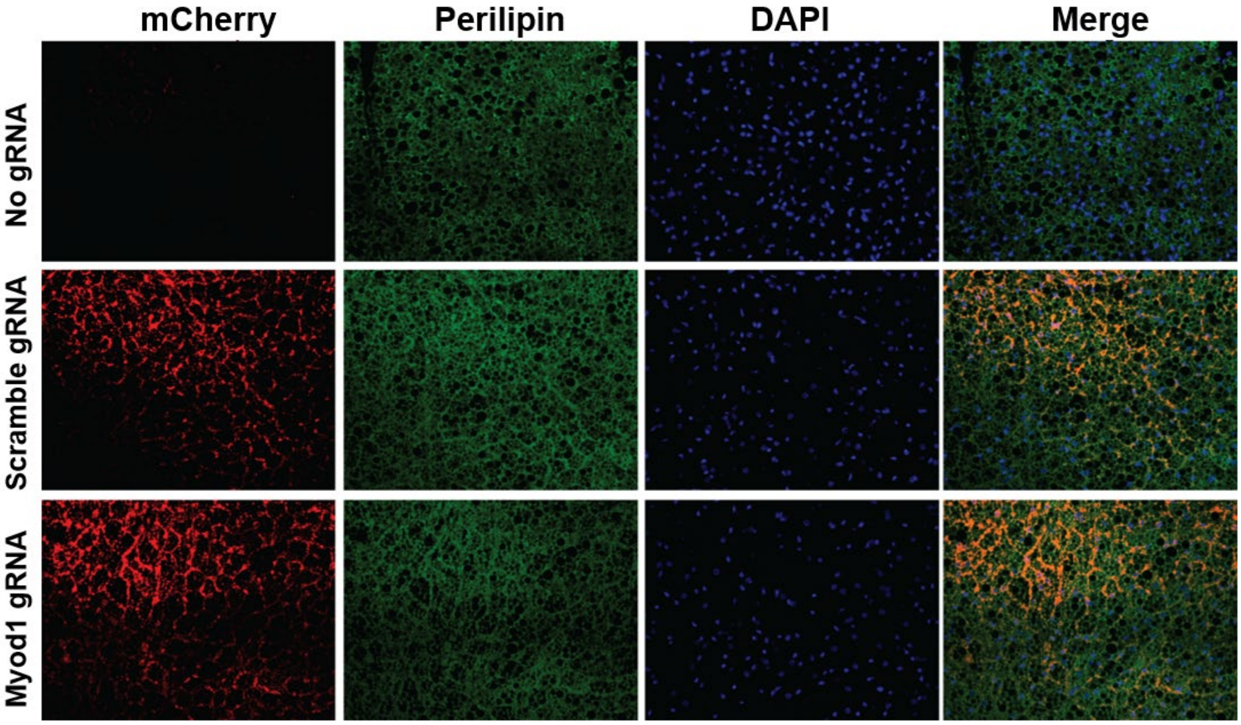


**Supplemental figure 16 (Related to Figure 5).** Representative images indicate IHC staining of mCherry (red), perilipin (green), DAPI (Blue) and merged images from iBAT of lentivirus-injected mice. Three-month-old chow-fed male C57BL/6J mice were bilaterally injected with lentiviruses expressing dCas9-TET1CD plus lentiviruses expressing either targeting Myod1-gRNA-mCherry or non-targeting scramble-gRNA-mCherry into iBAT. Tissues were collected 2 months after the injection.
